# A combined human *in silico* and CRISPR/Cas9-mediated *in vivo* zebrafish based approach for supporting gene target validation in early drug discovery

**DOI:** 10.1101/2021.09.14.460241

**Authors:** Matthew J. Winter, Yosuke Ono, Jonathan S. Ball, Anna Walentinsson, Erik Michaelsson, Anna Tochwin, Steffen Scholpp, Charles R. Tyler, Steve Rees, Malcolm J Hetheridge, Mohammad Bohlooly-Y

**Author notes:** **Corresponding authors:** Matthew J. Winter, Biosciences, College of Life and Environmental Sciences, University of Exeter, Geoffrey Pope Building, Stocker Rd, Exeter, Devon, EX4 4QD, United Kingdom. **Corresponding authors:** Mohammad Bohlooly, Translational Genomics, Discovery Sciences, BioPharmaceuticals R&D, AstraZeneca, Gothenburg, SE-431 83, Sweden. These authors have contributed equally to this work (Matthew J. Winter, Yosuke Ono, Jonathan S. Ball).

## Abstract

The clinical heterogeneity of heart failure has challenged our understanding of the underlying genetic mechanisms of this disease. In this respect, large-scale patient DNA sequencing studies have become an invaluable strategy for identifying potential genetic contributing factors. The complex aetiology of heart failure, however, also means that *in vivo* models are vital to understand the links between genetic perturbations and functional impacts. Traditional approaches (e.g. genetically-modified mice) are optimal for assessing small numbers of proposed target genes, but less practical when multiple targets are identified. The zebrafish, in contrast, offers great potential for higher throughput *in vivo* gene functional assessment to aid target prioritisation and support definitive studies undertaken in mice. Here we used whole-exome sequencing and bioinformatics on human patient data to identify 3 genes (*API5, HSPB7*, and *LMO2*) suggestively associated with heart failure that were also predicted to play a broader role in disease aetiology. The role of these genes in cardiovascular system development and function was then further investigated using *in vivo* CRISPR/Cas9-mediated gene mutation analysis in zebrafish. We observed multiple impacts in F0 knockout zebrafish embryos (crispants) following effective somatic mutation, including reductions in ventricle size, pericardial oedema, and chamber malformation. In the case of *lmo2*, there was also a significant impact on cardiovascular function as well as an expected reduction in erythropoiesis. The data generated from both the human *in silico* and zebrafish *in vivo* assessments undertaken supports roles for *API5, HSPB7*, and *LMO2* in human cardiovascular disease and identifies them as potential drug targets for further investigation. The data presented also supports the use of human *in silico* genetic variant analysis, in combination with zebrafish crispant phenotyping, as a powerful approach for assessing gene function as part of an integrated multi-level drug target validation strategy.

## Introduction

Chronic heart failure is characterised by a mismatch between cardiac output and the oxygen demands of organs. Behind the clinical syndrome there is a well-established sequence of pathophysiological events eventually resulting in maladaptive cardiac remodelling, ventricular dilatation and poor cardiac performance manifested as reduced ejection fraction (Konstam et al., 2011). Despite this, at least half of the heart failure population falls outside of the definition of heart failure associated with reduced ejection fraction (HFrEF), and although the aetiology of heart failure in HFpEF patients is largely unknown, a paradigm has been proposed arguing that the root cause is extracardiac (Senni et al., 2014). This multifactorial aetiology makes identifying potential drug targets for the treatment of chronic heart failure challenging. Target identification has been greatly aided by the emergence of large-scale patient DNA sequencing approaches (Suwinski et al., 2019; Povysil et al., 2020), although this strategy has its own limitations. Candidate gene target progression, for example, is often complicated by identification of multiple potential targets that require subsequent functional characterisation and prioritisation. This is compounded by the fact that higher throughput screening approaches are currently limited to *in silico* or *in vitro* methods that lack the ability to score organ system-dependent gene function. On the other hand, higher-tier genetic target validation assessments in traditional animal models are not practical for screening multiple candidate genes. As an alternative *in vivo* model, the embryo-larval zebrafish could fill this gap. The zebrafish combines genetic tractability, higher throughput amenability and optical transparency allowing the relatively simple assessment of organ system morphology and functionality across multiple gene targets (Gut et al., 2017). Importantly, the zebrafish is also widely considered to be an appropriate animal model for studying human cardiovascular biology (MacRae and Peterson, 2015). Furthermore, recent studies have demonstrated the great utility of zebrafish in CRISPR/Cas9 mediated screens using F0 knockouts (crispants) as rapid, highly reproducible and scalable knockout models (Burger et al., 2016; Kroll et al., 2021). Here, we used this approach to investigate the function of 3 genes implicated in human cardiovascular disease from a large-scale patient DNA sequencing study.

Whole exome sequencing (WES) and subsequent bioinformatics were used to identify genes from clinical cohorts that were suggestively associated with heart failure (Povysil et al., 2020). We identified a subset of three genes (*API5, HSPB7*, and *LMO2*) predicted to play a broader role in heart failure aetiology, and undertook *in vivo* phenotypic assessment in zebrafish. These specific genes were selected as they were representative of genes that: had broad pleiotropic functions without evidence of preferential cardiac expression (*API5*); showed preferential expression in the heart, but with ambiguous function (*HSPB7*); or were expressed in the haematopoietic compartment and thus had potential impacts on erythrocytes physiology, oxygen delivery and leukocyte biology (*LMO2*). The positive control gene selected was *GATA5*, which has a critical role in heart development and has been implicated in multiple human cardiovascular disease aetiologies (Gu et al., 2012; Wei et al., 2013; Zhang et al., 2015).

Functional knockout of *gata5* resulted in zebrafish larvae exhibiting the expected cardiovascular phenotype, and mutation of the three target genes resulted in some degree of negative impact on the physiology and/or development of the zebrafish cardiovascular system. The evidence presented supports the use of human *in silico* gene variant analysis in combination with zebrafish crispant assessment as a powerful strategy for assessing gene function during target identification and validation. Furthermore, the data generated provides strong *in vivo* evidence to support the further investigation of these genes as novel drug targets for treating human cardiovascular disease.

## Materials and Methods

### Case-control collapsing analysis

Target genes were identified by WES in heart failure patients from two clinical trials; Candesartan in Heart Failure-Assessment of Reduction in Mortality and Morbidity (CHARM) (Pfeffer et al., 2003) and Controlled Rosuvastatin Multinational Trial in Heart Failure (CORONA) (Kjekshus et al., 2007). 5942 heart failure cases from these trials were compared to controls without reported heart disease using gene-based rare-variant collapsing analysis (Povysil et al., 2020). One gene, TTN (encoding Titin), reached study-wide significance, with the strongest association in the dominant protein-truncating variant (PTV) model (p = 3.35×10^−13^), a finding that was replicated in the UK Biobank WES data (Povysil et al., 2020) and supported by our *in silico* analysis. From this, a list of 255 genes that had p-values above the study-wide significance threshold, but below 1 × 10^−4^, were further explored for data supporting a role in cardiovascular disease using a bioinformatics prioritisation assessment as described below.

### Bioinformatics analysis

Each gene was first assessed for genetic association to human disease phenotypes based on large-scale genome-wide association (GWAS) and WES studies encompassing common to low frequency variants (Common Metabolic Diseases Knowledge Portal or CMDKP, GWAS Catalog, Phenoscanner). Rare variant associations reported in Online Mendelian Inheritance in Man (OMIM) and ClinVar were also captured. Baseline tissue and cellular expression of candidate targets were investigated based on bulk and single cell RNA sequencing data from human tissues. Studies of expression dysregulation in cardiovascular disease were also conducted using patient transcriptomics data deposited in NCBI Gene Expression Omnibus (GEO) using QIAGEN’s OmicSoft DiseaseLand (release HumanDisease_B37_20191215 _v14a), which applies generalised linear models on log2 transformed intensities (microarray data), and DESeq2 for raw counts data (RNAseq data). Genes were considered significantly differentially expressed at an adjusted p<0.05. For mechanistic inference assessment, network-based functional enrichment analysis was performed using three separate tools (STRING, Harmonizome and GeneMANIA) all relying on multiple data types including protein-protein interactions, co-expression, database and text mining. From these *in silico* analyses, *API5, HSPB7* and *LMO2* were selected for *in vivo* assessment.

### Guide RNA (gRNA) design and preparation

For the zebrafish orthologues of each target gene (*gata5∼*ENSDARG00000017821; *api5∼*ENSDARG00000033597; *hspb7∼*ENSDARG00000104441; *lmo2∼*ENSDARG00000095019), three individual guide RNAs (gRNAs) were designed (https://chopchop.cbu.uib.no) to target discrete sections of coding exon 1, except for *lmo*2 which targeted exon 2 (**Fig. S1**). Each gRNA was applied alone (termed g#1, g#2 or g#3) to assess the consistency of phenotypes across different target sites, and as a combined injection containing all three gRNAs (termed g#1,2,3) to ensure functionally-effective mutation, alongside Cas9-only injection controls. Prior to injection, gene-specific crRNA and tracrRNA (Integrated DNA Technologies Inc. Coralville, USA) were diluted to a final concentration of 12 μM in nuclease-free duplex buffer and the resultant gRNA mixture incubated at 95°C for 5 mins. Immediately prior to use, 5 μl of the gRNA mixture was mixed with Cas9-NLS protein (final concentration of 5 μM. New England Biolabs, Ipswich, USA), 2M KCl (final concentration of 300mM), and 0.5% v/v Phenol red solution (Sigma Aldrich UK Ltd. Poole, UK). The resultant mixture was incubated at 37 °C for 10 mins to assemble the gRNA/Cas9 ribonucleoprotein complex, and then held at room temperature until use.

### Zebrafish culture

Adult WIK (Wild-type India Kolkata) strain zebrafish (*Danio rerio*), originally obtained from the Zebrafish International Resource Center (ZIRC, University of Oregon, Eugene, USA), were held under optimal spawning conditions (14 hour light: 10 hour dark cycle, with 20 minute dusk-dawn transition periods, 28 ± 1°C), in mixed sex groups in flow through aquaria. Each injection day embryos were collected from individual male-female pairs and injected at the one-cell stage. In addition, the *cmlc2::DsRed2-nuc* transgenic zebrafish (Mably et al., 2003) used for confocal cardiomyocyte microscopic analysis were cultured under identical conditions.

### Microinjection

Microinjection needles were prepared from thin wall borosilicate glass capillaries with filament (Outer diameter 1.0mm, inner diameter 0.75mm. World Precision Instruments, Sarasota, USA) on a micropipette puller (P-1000,Sutter Instruments, Novato, USA) using the following settings: Heat 501, Pull 60, Velocity 60, Time 20, Pressure 300, Ramp 499.

From pairs of spawning zebrafish, eggs were assessed for the desired development stage (1-cell) and for condition before being transferred in batches of 50-60 into the furrows of an injection mould-imprinted agar plate. Next, the injection needle was loaded with the injection mixture calibrated using a microscale graticule to deliver 1-1.7nl per injection and each egg was injected (FemtoJet 4x, Eppendorf, Hamburg, Germany), once, close to the cell/yolk boundary layer. Successful injection was indicated by the presence of phenol red. Injected eggs were then transferred to a Petri dish containing culture water and methylene blue (2 drops per litre of water) and cultured on a black background under the same conditions as the adult fish. At the end of day 0, all unfertilised and dead embryos were removed and 48 viable embryos, selected at random for each treatment, were individually transferred to wells of 48-well microplates (each in 1ml) for later assessment.

### Morphological assessment at 2 and 4 dpf

At 2 days post fertilization (dpf), the general morphological phenotype of all embryos across all 3 individual and a combined gRNA injected groups (named g#1, g#2, g#3 and g#1,2,3 respectively) was assessed to identify the most effective guides (i.e. those resulting in the most prominent phenotype versus the Cas9-only injected controls) for complete morphological and functional phenotyping at 4 dpf. Scoring was undertaken (without anaesthesia) using a dissecting microscope against a list of criteria shown in **supplementary Table S2**. In addition, 8 embryos were removed from each treatment for analysis of gene mutation efficiency (**see below**).

At 3 dpf, if necessary, embryos were manually dechorionated using fine forceps allowing the spine to straighten to facilitate complete phenotyping at 4 dpf. At 4 dpf, 10 animals were selected at random from the 2 most effective treatments groups, alongside 10 embryos from the Cas9-only injected control group for full morphological scoring using a method based upon Gustafson *et al*., (Gustafson et al., 2012) and Ball *et al*. (Ball et al., 2014). To facilitate scoring, animals were lightly anaesthetised by immersion in 0.165 g/L tricaine methanesulfonate (pH 7.5) and scored according to the criteria shown in **supplementary Table S3**. Images were taken from representative animals within each treatment group. After scoring, each animal was directly transferred to benzocaine solution (1 g/L in 1% ethanol) for euthanasia. The guide-injected group providing the most robust phenotype was also selected for a second run to confirm the observed effect in a separate batch of embryos.

### Analysis of mutation efficiency

Genomic DNA was extracted from individual 2 dpf larvae using the HotSHOT method (Meeker et al., 2007). Briefly, all water was removed from each PCR tube containing an embryo, 50 μl of 50 mM NaOH added, and the sample heated for 10 mins at 95°C. The samples were then vigorously vortexed and subsequently cooled on ice. Next 5 μl of 1M Tris-HCl (pH 8.0) was added and the samples were well mixed. The samples were then centrifuged and the supernatant containing the genomic DNA was removed and stored at -20 °C until further processing. The PCR primers were designed and obtained from Eurofins Genomics (Ebersberg, Germany). The primers used were: *gata5* (forward: GGAAACCATCGCATTTGGAG and reverse: AGGGCACTTCCATATTGATC); *api5* (forward: ATACAGCGGAAGTATCCGAC and reverse: TCAATTCTCGCTCAGGCTTG; *hspb7* (forward: GAATAAGAACTTGATCACCGG and reverse: GCATATAGCTTTCCACTCAC); and *lmo2* (Forward: TGGATGAGGTGCTCCAGATG and reverse: ATCTCTCCTGCACAGCTTTC). The PCR mixture was prepared as follows: 10 μl of 2x PCRBIO Taq Mix Red (PCR Biosystems Ltd., London, UK); 0.8 μl each of 10 μM forward and reverse primers; 1 μl of genomic DNA; and 7.4 μl of water. The PCR machine settings were as follows: 1 min at 95°C; 30 cycles of 15sec at 95°C, 15sec at 58°C, 15sec at 72°C; and finally 1 min at 72°C. DNA amplification was checked on a 3% agar gel. T7 endonuclease I (T7E1) assays were undertaken to detect heteroduplexes in the PCR product. For this, PCR products were denatured at 95°C for 5 mins and then cooled down. Next, the T7E1 reaction mixture was made as follows: 10 μl of each PCR product; 1.5 μl NEBuffer 2 (New England Biolabs, Ipswich, USA); 0.3 μl T7E1 enzyme at 10 Units/μl (New England Biolabs, Ipswich, USA); and 3.2 μl water. Next digestion was undertaken for 15 min at 37°C and the resultant products were assessed on a 3% agar gel. A sub-sample of amplified DNA from each PCR product was also sent for Sanger sequencing by Eurofins Genomics (Ebersberg, Germany) using the forward or reverse primers described previously.

### Histology

For histological analysis, 4 dpf animals were terminated by anaesthetic overdose in fixation tubes (2 g/L tricaine methanesulfonate, pH 7.5). Next the anaesthetic was replaced with 10% neutral buffered formalin for 4 hours, followed by 70% ethanol/methanol in which they were stored at 4°C until further processing. For sectioning, animals were transferred into agar moulds for orientation (Sabaliauskas et al., 2006; Copper et al., 2018), and subsequently into tissue cassettes and embedded in paraffin using an automatic tissue processor (Thermo Scientific Excelsior AS, Thermo Fisher Scientific Ltd., Waltham, USA). The sequence of fixation and embedding steps applied are summarised in **Supplementary Table S4**. Following fixation, 5 μm sections were cut from each paraffin block using a microtome (Shandon AS325, ThermoFisher Scientific, Waltham, USA). The resultant sections were haemoxylin and eosin (H&E) stained on an automatic stainer (Shandon Varistain 24-4, ThermoFisher Scientific, Waltham, USA) using the sequence summarised in **Supplementary Table S5**. After staining, images of each section were captured on a binocular microscope (Axioskop 40, Zeiss, Oberkochen, Germany) equipped with a colour digital camera (DP70, Olympus, Tokyo, Japan) to allow histopathological analysis.

### Cardiovascular functional assessment at 4 dpf

Cardiovascular function was assessed in ten 4 dpf embryos selected at random from each treatment, using the method outlined in Parker *et al*. (Parker et al., 2014). Briefly, each animal was lightly anaesthetised by immersion in tricaine methanesulfonate (0.1 g/L pH 7.5) and then transferred into low melting point agarose (1 g/100 ml of the same tricaine methanesulfonate solution to maintain anaesthesia during imaging) and then deposited on its side on a clear microscope slide. Imaging was undertaken on an inverted light microscope (Leica DM IRB, Leica Microsystems UK Ltd., Milton Keynes, UK, 10x magnification) equipped with two video cameras: one recording the heart at 25 frames per second (fps. Grasshopper® GRAS-50S5C-C, Point Grey, Richmond, Canada); and the second recording the dorsal aorta at 120fps (Grasshopper® GRAS-03K2M-C, Point Grey, Richmond, Canada). Recording was undertaken for 10 mins following which animals were directly transferred to benzocaine solution (1 g/L in 1% ethanol) for euthanasia without recovery.

Heart videos were analysed using MicroZebraLab™ (v3.5, ViewPoint, Lyon, France) from which beat frequencies were provided for each of the atrium (atrial beat rate or ABR) and the ventricle (ventricular beat rate or VBR). This allowed global heart rate measurement and the detection of certain arrhythmias such as the decoupling of atrial and ventricular beat frequencies, which has been previously described in association with exposure to some QT-prolonging drugs. Blood flow videos were analysed using ZebraBlood™ (v1.3.2, ViewPoint, Lyon, France), which provided measures of blood flow (nL/sec), blood linear velocity (μm/sec) and vessel diameter (μm). In addition to the directly determined parameters, estimates of stroke volume and cardiac output were calculated using measurements of heart rate and blood flow (termed surrogate stroke volume (SSV) and surrogate cardiac output (SCO)). Normally stroke volume is precisely calculated using the difference between end-systolic and end-diastolic volumes (see below). However, using our system a surrogate measure of SSV was calculated by dividing the dorsal aorta flow rate (in nL/sec), by the VBR per second. Similarly, cardiac output is normally calculated by dividing the stroke volume by the heart rate to provide a volume of blood pumped per minute. Here, however, SCO was calculated by multiplying the SSV by the VBR in bpm.

### Ventricular dimension-related parameter measurements

Using the videos of the heart captured for functional analyses, 10 animals were randomly selected per treatment (5 per run) from which a manual measurement of ventricle diameters was undertaken. Using VirtualDub (http://www.virtualdub.org/), heart videos were converted to JPEGs from which 10 images showing the minimum (end-systolic) and 10 showing the maximum (end-diastolic) ventricle chamber extension were selected at random from each animal. On each image, the long and short axis ventricle chamber lengths were measured using ImageJ (https://imagej.net/). Next, using the equation 1/6*π*long axis*short axis^2^ and assuming a prolate spheroidal shape (Yalcin et al., 2017), end-systolic and end-diastolic ventricle volumes were calculated (in nL). From these measurements, stroke volume (end-diastolic minus end-systolic volumes, in nL), cardiac output (stroke volume*ventricular beat rate, in nL/min), and ejection fraction (stroke volume/end-diastolic volume*100, as a %) were calculated to supplement the surrogate measures of cardiac performance described previously. Although of much slower throughput, manual measurement of ventricular diameters is particularly useful where the absence of measurable blood flow means that SSV and SCO values are effectively zero. It should be noted, however, that as 2D images were derived from videos that were used for the functional analysis, in some cases precise determination of chamber edges was difficult and as such the actual dimensions should be considered approximations. In addition, it was not possible to normalise these measurements to the length of the animal used for functional assessment. However, ventricle measurements have been considered within the context of the average total body length of embryos within the same batch of animals used for the morphological evaluation.

### Confocal analysis of cardiomyocyte development and morphology

To assess the impact of gene mutation on cardiomyocyte hyperplasia or hypertrophy, the mutation of each target gene was analysed visually in 4 dpf *cmlc2::DsRed2-nuc* transgenic zebrafish. For each gene and for Cas9-only injected animals, 10 randomly selected animals were assessed on a Nikon A1R laser scanning confocal microscope (Nikon, Tokyo, Japan) using 568nm laser excitation (power 90, PMT 85) and transmitted light (PMT 25). At 20x magnification, Z-slices were taken every 5 μm through the heart from which maximum intensity z-projections were then generated. For imaging, the embryo’s hearts were stopped by immersion in 1 g/L tricaine methanesulfonate (pH 7.5) after which they were transferred to 1 g/100 ml low melting point agarose made using the same tricaine methanesulfonate solution for immobilization during imaging. As before, after imaging larvae were directly transferred to a solution of benzocaine (1 g/L in 1% ethanol) for euthanasia without recovery. Each maximum intensity projection was imported into ImageJ, and the brightness was adjusted to aid visualization of cardiomyocyte nuclei. From these images, cardiomyocyte number was estimated in the ventricle of each animal using a manual cell counter.

### Data analysis

All measured parameters were averaged per animal to provide a series of individual values from which the treatment mean and SEM were calculated. For statistical analysis, each group was first tested for normality (Anderson-Darling Test) and homogeneity of variance (Levene’s, Bartlett’s, or F-test). Each treatment was then compared with the Cas9-injected control group using either the Student’s T-tests or 1-way ANOVA and Tukey’s HSD tests (parametric), or the Mann Whitney U-tests or Kruskal Wallis and Dunn’s Tests (non-parametric). All analyses were undertaken using Minitab™. Throughout data are shown as the mean, ± standard error of the mean (n), with a minimum *α* level of 0.05 applied (with a Bonferroni correction in the case of multiple comparisons).

### Animal Ethics statement

All animal work was carried out in accordance with the EU Directive for the protection of animals used for scientific purposes (2010/63/EU) and UK Animals Scientific Procedures Act (ASPA) 1986. Experimental procedures were carried out under personal and project licenses granted by the UK Home Office under ASPA, and ethically approved by the Animal Welfare and Ethical Review Body at the University of Exeter.

## Results

From WES and subsequent bioinformatics, three genes were identified as suggestively associated with heart failure and were considered representative of groups of genes predicted to play a broader role in heart failure aetiology. These genes were then subjected to *in vivo* phenotypic assessment in zebrafish alongside the positive control gene *GATA5*, in order to further investigate their role in vertebrate cardiovascular development and function.

### *In silico* analysis of clinical data

Key results of the *in silico* analysis are summarised in **Fig. 1**. Additional data are contained within **Supplementary figures S2-S3 and supplementary data S1-S4**, and literature-derived information summarised in **Supplementary Table S1**.

**Fig. 1.**
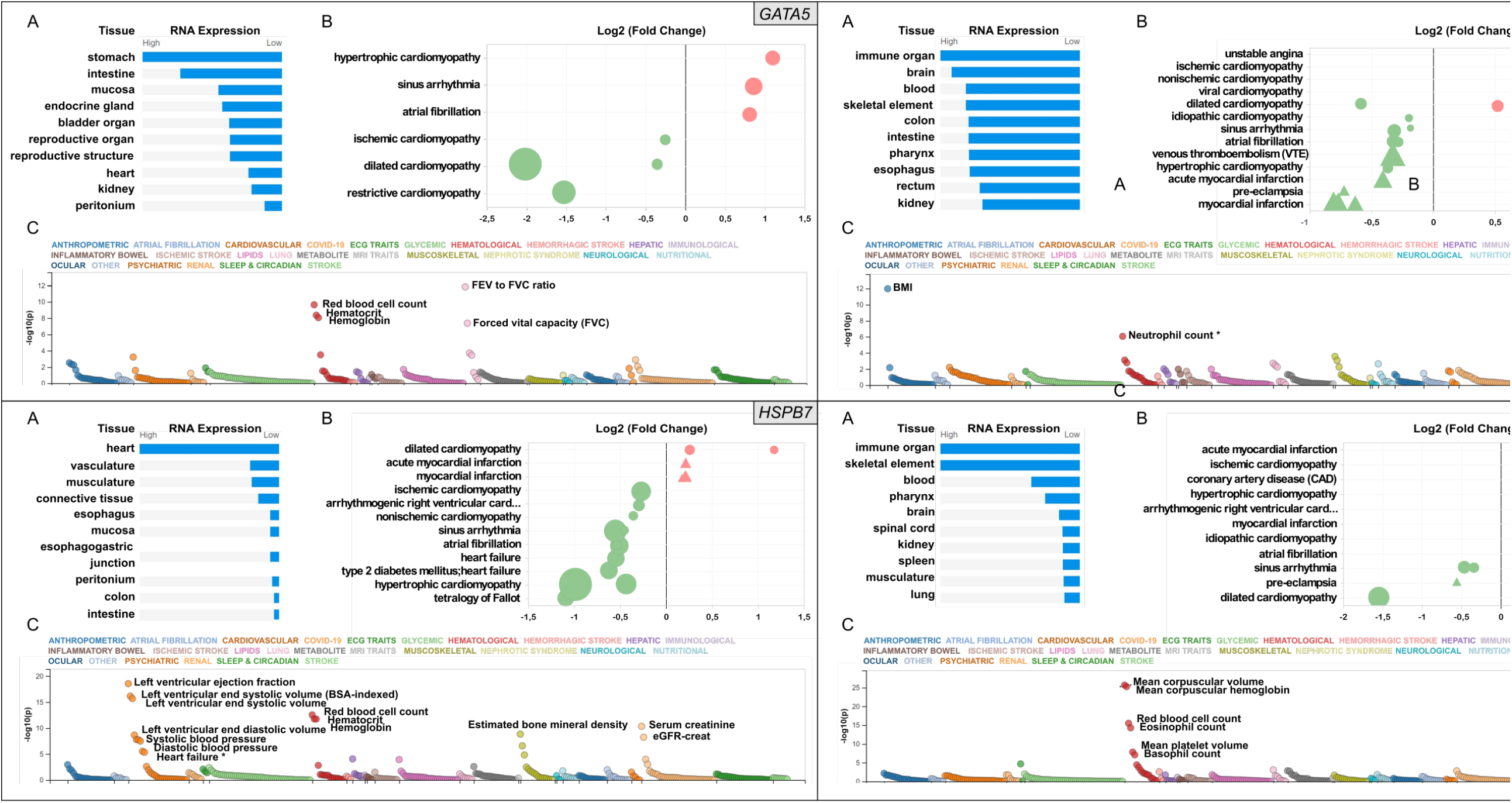
Snapshot of the results from the *in silico* assessment of the heart failure candidate genes *API5* (top right), *HSPB7* (bottom left), *LMO2* (bottom right), and positive control gene *GATA5* (top left panel). **A**. The top 10 tissues showing expression based on RNA sequencing data from the Human Protein Atlas (HPA) and EMBL-EBI Expression Atlas as summarized by Open Targets Platform (https://www.targetvalidation.org). **B**. Gene expression changes in cardiovascular disease conditions based on publicly available transcriptome studies from NCBI Gene Expression Omnibus (GEO). Disease conditions are shown on the Y-axis and log2 fold changes (vs normal controls) on the X-axis. Icons are coloured by direction of change; red and green represent up- and downregulation in that disease, respectively. Icon shapes represent tissue type subjected to transcriptomics; circles and triangles represent heart and blood, respectively. Finally, icon size reflects statistical significance; the larger the icon the lower the P-value. All findings shown are significant (Adjusted P-value<0.05). **C**. Common variant gene locus association data from 190 datasets and 251 traits in Common Metabolic Diseases Knowledge Portal (CMDKP). Traits considered genome-wide significant (p-value ≤ 5×10^−8^) are highlighted (border-line significant traits are marked with *).

Relative tissue mRNA expression levels of *GATA5* are shown in **Fig 1A**. As expected, heart *GATA5* mRNA expression (**Fig. 1B)** was significantly altered in association with various cardiovascular disease states, the most significant being downregulated expression in dilated cardiomyopathy. The most pronounced changes in cardiac and blood *API5* expression included downregulated expression in myocardial infarction, and upregulated heart *API5* expression in ischemic and non-ischemic cardiomyopathies. Heart *HSPB7* expression was largely downregulated in various cardiac disease conditions, except for dilated cardiomyopathy in which *HSPB7* was found to be upregulated in two independent studies. *LMO2* blood mRNA expression predominantly showed upregulation, including in association with ischemic cardiomyopathy, coronary artery disease and myocardial infarction. In contrast, cardiac *LMO2* mRNA expression showed both increases and decreases across the same range of conditions.

Analysis of common variant gene-level associations (**Fig. 1C**) revealed that *GATA5* was significantly associated with various lung functions and haematological traits. Despite this, no significant common variant associations with heart disease were found, although rare loss-of-function mutations in *GATA5* have been reported to cause congenital heart defects (Jiang et al., 2013). Assessment of *API5* revealed a significant association with body mass index (BMI) and a borderline significant association with neutrophil count (p-value=9 × 10^−7^). Among the cardiovascular traits assessed, low frequency 3’UTR or intron *API5* genetic variants showed a significant association with “Cause of death: atrial fibrillation and flutter” (p=6.4^-22^), and “Cause of death: acute and subacute infective endocarditis” (p=6.7^-13^). *HSPB7* common variant data showed a significant association with cardiovascular, renal, haematological and musculoskeletal traits with the most significant association to cardiac function including left ventricular ejection fraction, and left ventricular end-systolic volume. Other GWAS data revealed significant associations between common variants in *HSPB7* intron 3’ and 5’UTR and idiopathic dilated and sporadic cardiomyopathy (p=5.3^-13^ and p=1.4^-9^ respectively), as well as with systolic blood pressure (p=7^-12^). Finally, analysis of the *LMO2* data revealed significant associations with various haematological traits including mean corpuscular volume, haemoglobin and red blood cell count. The most significant cardiovascular trait association was only suggestive (P-wave duration at p=2.1^-5)^. Further analysis did, however, reveal that *LMO2* intron variants were significantly associated with “Cause of death: cardiomegaly” (p=3.0^-9^) and “Cause of death: dilated cardiomyopathy” (p=1.3^-8^).

### *In vivo* target gene mutation efficiency

Site-specific mutagenesis in zebrafish embryos were evaluated by T7E1 assay and sequencing of genomic PCR products amplifying the region that gRNAs target. There was clear cleavage of PCR products by T7E1 in the presence of Cas9 and gRNAs, demonstrating highly efficient mutagenesis across the four targets (**Fig. 2**). In addition, sequencing confirmed that effective site-specific gene mutation was achieved for all combinations of gRNAs, but not in the case of the Cas9-only injected controls (**Fig. 2**).

**Fig. 2.**
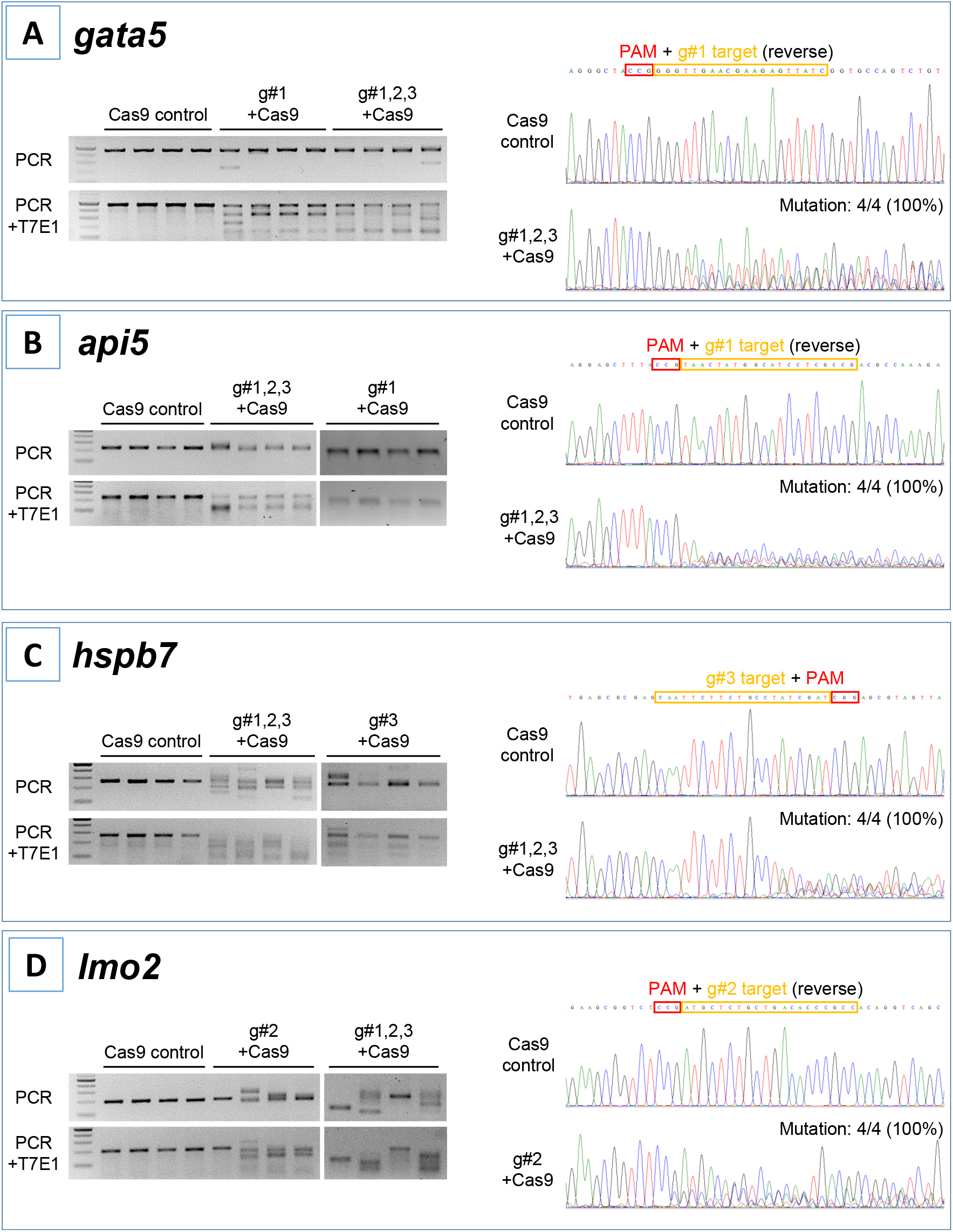
Mutation efficiency of the gRNAs for each target gene. For each gene, the left-hand gel images show (upper image) the bands obtained following targeted PCR of genomic DNA extracted from 4 individual animals injected with the two most effective CRISPR gRNAs + Cas9 (based on 2 dpf morphological analysis), compared with the Cas9 injected control animals. The lower gel images show the same samples following T7E1 assay undertaken to reveal the cleavage of heteroduplex DNA. The right-hand image in each case shows the result of Sanger sequencing undertaken on representative genomic DNA samples from the most effective gRNA + Cas9, per target gene, compared with that from a representative Cas9-injected control animal (top image). Data are shown for A) *gata5*, the positive control gene, B) *api5*, C) *hspb7* and D) *lmo2*. Note in all cases the most effective gRNA was the combined guide group (g#1,2,3) except for *lmo2* which due to high mortality of the g#1,2,3 group, the g#2-injected animals were selected for full analysis. On the right-hand edge of each panel, the number of animals that showed mutation in the target region by sequencing is shown. In all cases 4/4 (100%) showed mutation in the presence of gRNAs + Cas9, but zero in the Cas9-only control group animals.

### Morphology and function of crispants

At 2 dpf, the most robust *gata5* crispant phenotypes occurred after injection of g#1,2,3, followed by injection of g#1 alone (**Fig. S4**). By 4 dpf (**Fig. 3**), all *gata5* crispants showed pericardial oedema, and there were high incidences of misshapen and small heart chambers (e.g. 75% of the combined gRNA group) together with a frequent lack of chamber definition (e.g. 40% of the combined gRNA group). In addition, both groups of *gata5* crispants exhibited a range of non-cardiac developmental abnormalities (**Fig. 3**) that included an increased incidence of poorly defined somites, malformed fins, small and malformed eyes, reduced neural tube size, a lack of definition of the fore-midbrain boundary and reduced size of the olfactory region. In addition, the g#1,2,3 group exhibited a comparatively high incidence of malformed brachial arches and deficient or absent jaw structures. Histological analysis of the heart (**Fig. 4A and Fig. S5**) revealed that *gata5* crispants generally exhibited cardiac hypoplasia, chamber malformation, an absence of visible heart valves and pericardial distension. This abnormal cardiac phenotype was further supported after the analysis of ventricle dimensional parameters (**Fig. 4B**), which revealed significantly smaller end-diastolic and end-systolic ventricle diameters and volumes, as well as reduced stroke volume, cardiac output and ejection fraction in the *gata5* crispants versus the Cas9 control animals. Furthermore, *gata5* crispants (**Fig. 4C and Fig. S6**) exhibited weaker and patchy DsRed2 fluorescence, reduced numbers of cardiomyocytes (estimated at 18 ±4.2 versus 70 ±3.06 in the Cas9 controls. Mean, ± SEM, n=10) and predominantly misshapen and small cardiomyocyte nuclei. The abnormal cardiac phenotype was reflected in reduced cardiovascular function in the *gata5* crispants. For all endpoints measured except vessel diameter (only measurable where blood flow occurred), *gata5* crispants showed reduced cardiovascular function compared with the Cas9 controls (**Fig. 5**, with accompanying videos in **Supplementary Videos**). Collectively these data supported a negative impact of *gata5* mutation on 4 dpf zebrafish cardiovascular physiology, supporting the validity of our screening approach in zebrafish crispants.

**Fig. 3.**
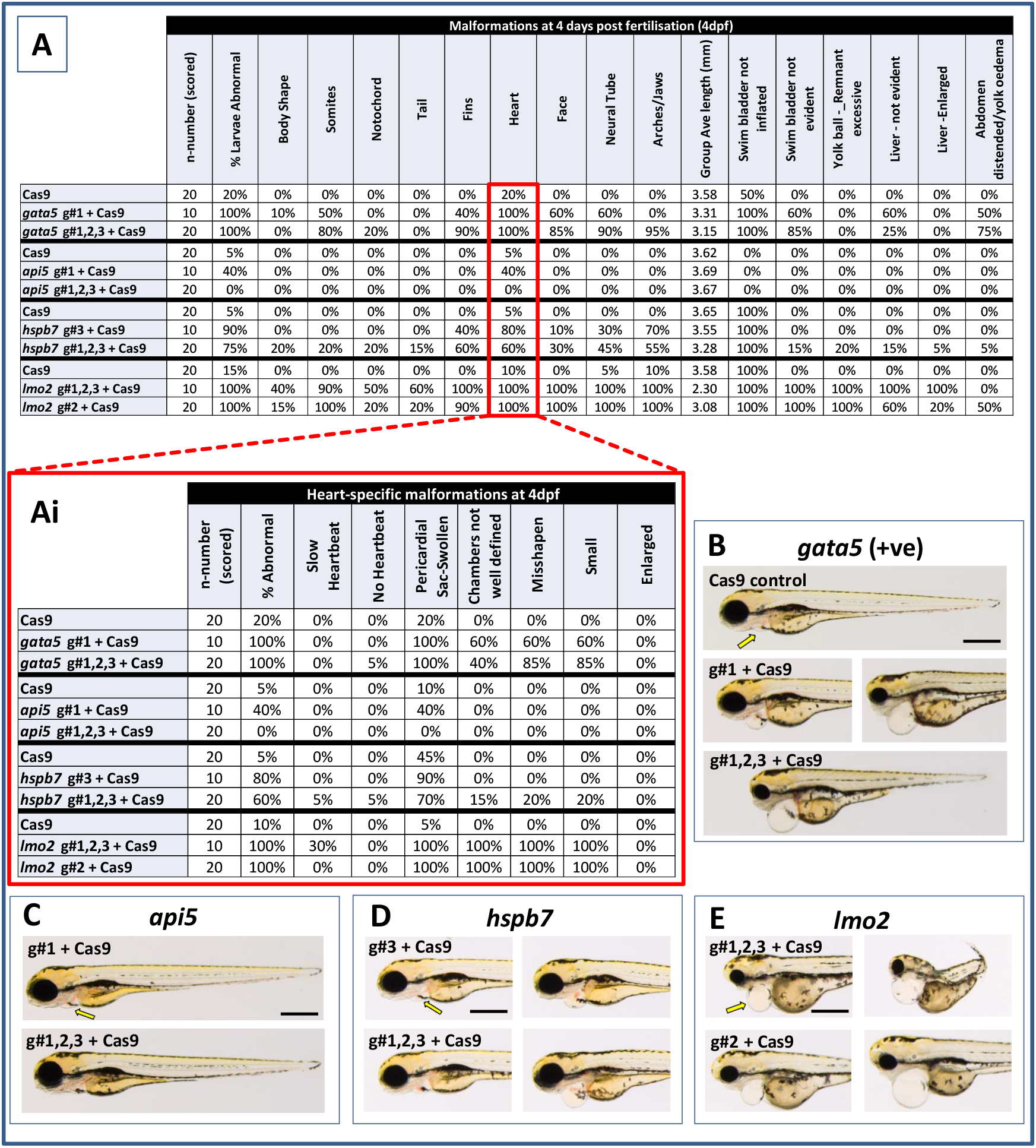
Results of the morphological analysis of 4 dpf *gata5* (positive control), *api5, hspb7* and *lmo2* zebrafish crispants versus the Cas9-injected control animals. **Panel A**: General whole body morphological endpoints measured following injection of Cas9 alone, or after mutation of each of the target genes. Data are shown as the % incidence of abnormalities under each category. Note different n-numbers present as two runs were undertaken for the Cas9 control and the gRNA + Cas9-injected group showing the most robust phenotype from run 1 (for *lmo2* g#2 was run twice due to concerns about excessive mortality in the g#1,2,3 group). The guide combination used for two runs in each case is shown in the lower panel of the example images for each gene. **Ai**: Expansion of heart-specific endpoints showing the full range scored. **B**: example larvae following *gata5* mutation versus the Cas9-injected control. The yellow arrows indicate the position of the pericardial membrane and the extent of pericardial oedema, which was minimal in the controls but extensive in most crispant animals (two examples are shown for g#1-injected animals as there was some variability in the severities seen). **C-E**: examples of larvae from other target gene treatment categories (note the apparent lack of effect of *api5* mutation on general morphology). The scale bar shown in the first image of panels B-E represents 500 μm.

**Fig. 4.**
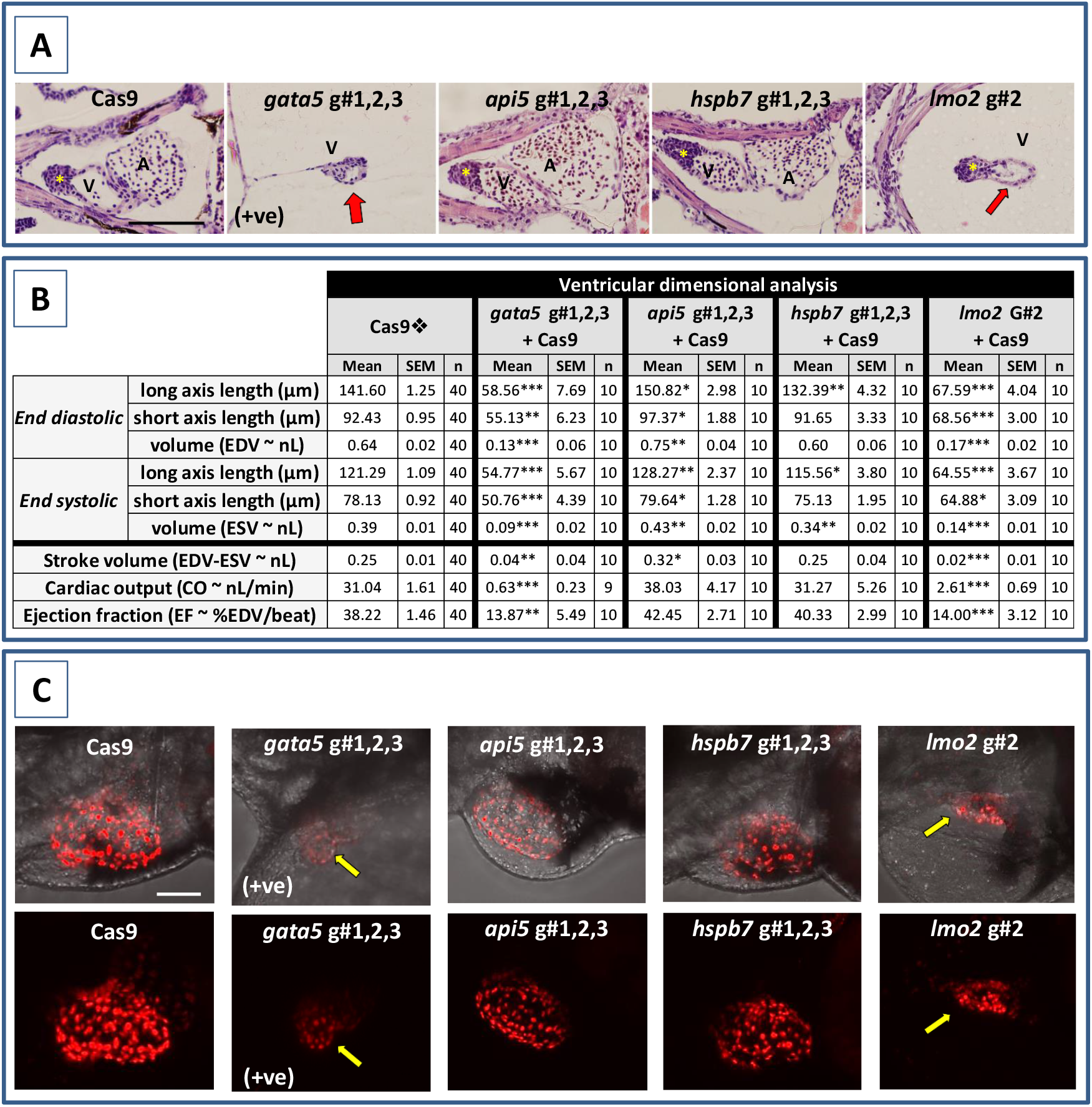
Results of the cardiac pathological analysis of 4 dpf *gata5* (positive control), *api5, hspb7* and *lmo2* zebrafish crispants versus the Cas9-injected control animals. **Panel A)** Example haematoxylin and eosin stained coronal sections through the heart (top, A = atrium, V = ventricle, *bulbous arteriosus) from each of the treatment groups versus the Cas9-injected controls (left-hand panels). Note in particular the extreme cardiac hypoplasia after *gata5* and *lmo2* mutation (indicated by a red arrow in the images) in which the atrium is not visible probably due to the severe pericardial oedema and resultant distension of the heart muscle. In each panel, animals are orientated with the head to the left, and viewed in the dorsal plane at a magnification of 40x (the scale bar shown in right-hand panel represents 200 μm). **Panel B)** Results of the analysis of ventricular dimensional analysis of the crispant versus Cas9 control animals. Shown are the ventricle dimensions and cardiac functional parameters calculated from the measurement of ventricle dimensions in 5 randomly selected embryos from each of the two runs undertaken on each gene. Data are shown as the mean, ± SEM, and n=animals. 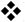 For brevity the Cas9 data are shown as the mean across all 4 Cas9 datasets (n=40 animals). However, statistical analysis was undertaken on the crispants versus the corresponding Cas9 control data in each case. *signifies a statistically significant difference versus the Cas9 control for that parameter at p<0.05, ** at p<0.01, and *** at p<0.001 (Student’s T-test or Mann Whitney U-tests). **Note**: the overall body lengths of the *gata5* and the *lmo2* crispants were also significantly reduced (p<0.001). **Panel C)** Example images of hearts from *cmlc2::DsRed2-nuc* larvae in which the cardiomyocytes are labelled red, especially prominently in the ventricle. The top row of panels shows the image with transmitted light and *cmlc2::DsRed2-nuc* fluorescence signals, and the lower row shows the same example but with the *cmlc2::DsRed2-nuc* signal alone. Note the severe oedema, weaker *cmlc2::DsRed2-nuc* fluorescence signal, reduced number of cells and smaller chamber size typical of the *gata5* crispant (indicated by the yellow arrow in images); the oedema, and slightly enlarged ventricle observed in the *api5* crispant; and the severe oedema, disorganisation of myocytes and smaller chamber size typical of the *lmo2* crispants (yellow arrow). *hspb7* crispant larval hearts outwardly appeared no different to the Cas9 controls. The scale bar shown in the upper left-hand image represents 50 μm.

**Fig. 5.**
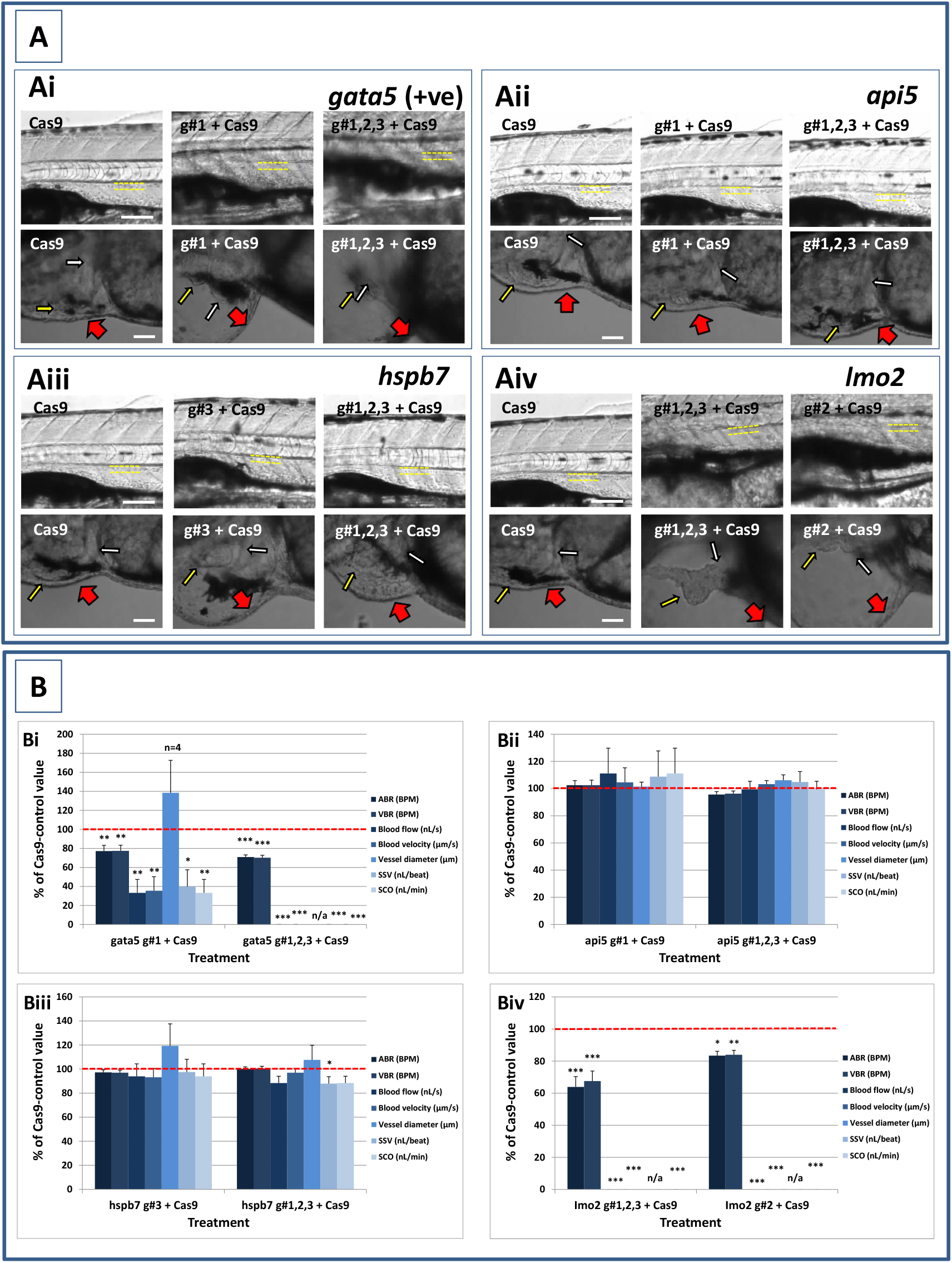
Results of the analysis of cardiovascular function in 4 dpf *gata5* (positive control), *api5, hspb7* and *lmo2* zebrafish crispants versus the Cas9-injected control animals. **Ai-Aiv:** Images of example Cas9 control larvae alongside larvae treated with the two gRNAs + Cas9 mixtures giving the most robust phenotypes (as assessed at 2 dpf). The top row shows the trunk vasculature with the position of the dorsal aorta outlined in yellow dashed lines, where blood flow and vessel diameter measurements were taken. The lower row of images shows the heart from the same animals, with the atrium highlighted by a small white arrow, and the ventricle by a small yellow arrow. The large red arrows show the position of the pericardial membrane and the extent of pericardial oedema. Most Cas9 control animals exhibited normal morphology and function in contrast with many of the crispants. **Bi-Biv:** cardiovascular functional endpoints quantified in the same groups of animals. Note the complete absence of blood flow measured in all of the *gata5* g#1,2,3- and in 6/10 of the g#1-injected animals, and the absence of effective blood flow in the *lmo2* g#1,2,3 and g#2-injected animals due to the absence of erythrocytes, meaning flow was not visible. Vessel diameter measurements were not possible in animals lacking blood flow (indicated by n/a). Data are shown as the mean % increase versus the Cas9-control group (100% indicated by the red dashed line), ± SEM, n=19-20 for the Cas9 and right-hand crispant treatment for each gene (data combined from two runs) and 10 for the left-hand treatment group for each gene where only one run was undertaken. *signifies a statistically significant difference versus the Cas9 control at p<0.05, ** at p<0.01, and *** at p<0.001 (T-test or Mann Whitney U-tests for the combined guide injected groups, or 1-way ANOVA and Tukey’s HSD tests or Kruskal-Wallis and Dunn’s tests for the single guide injected groups in which runs 1 and 2 were combined). The scale bar shown in the upper left-hand image of each panel represents 100 μm. The scale bar shown in the lower left-hand image of each panel represents 50 μm as a higher magnification camera mount was used in this case.

At 2 dpf the most robust *api5* crispant phenotypes were observed after injection of g#1,2,3, followed by g#1 alone (**Fig. S4**), although by 4 dpf there was little indication of any gross morphological impact other than a 40% incidence of very mild pericardial oedema in the g#1-injected animals (**Fig. 3**). Histology (**Fig. 4A and Fig. S5**) suggested a slight enlargement of the heart chambers with myocardial wall thinning, although this varied between individual animals. Chamber enlargement was, however supported by a small but significant increase in end-diastolic and end-systolic ventricle dimensions and volume in the *api5* crispants (**Fig. 4B**), which also resulted in a small, but significant, increase in stroke volume (although this was not evident from the video tracking-based analysis of cardiovascular function. See below). Confocal assessment of *cmlc2::DsRed2-nuc* animals (**Fig. 4C and Fig. S6**) revealed that *api5* crispants exhibited no evidence of an impact on cardiomyocyte number (estimated at 73 ±4.2 versus 70 ±3.06 in the Cas9 controls. Mean, ± SEM, n=10) and no obvious abnormal cardiomyocyte organisation or ultrastructure was observed. The apparent mild impact of *api5* mutation was also reflected in the absence of any significant effects on cardiovascular function after video tracking-based assessment (**Fig. 5 and Supplementary Videos**).

The most robust *hspb7* crispant phenotypes at 2 dpf occurred after injection of g#1,2,3, followed by g#3 alone (**Fig. S4**), and at 4 dpf there were widespread developmental abnormalities across multiple tissues compared with the Cas9 controls (**Fig. 3**). These abnormalities were particularly prevalent in the g#1,2,3-injected animals and included a 60% incidence of bent (predominantly pectoral) fins, a 45% incidence of a compressed/reduced size or malformed forebrain, and a 70% occurrence of malformed branchial arches or upper and lower jaw structures. In addition, there was a high incidence of pericardial oedema (60-80% of crispants) and in 20% of g#1,2,3 injected animal, misshapen, poorly-defined and small heart chambers. Histology (**Fig. 4A and Fig. S5**), however, did not reveal any clear ultrastructural abnormalities. A significant reduction in the end-diastolic and end-systolic long axis ventricle diameters was, however, detected along with a significant reduction in end-systolic volume suggesting a reduction in ventricle size (**Fig. 4B**). Analysis of cardiomyocyte organisation in *cmlc2::DsRed2-nuc* animals (**Fig. 4C**) also revealed some evidence of disorganised distribution of cardiomyocytes in the myocardium, and a small decrease in the numbers of cells present after *hspb7* mutation (estimated at 64 ±3.06 versus 79 ±2.78 in the Cas9 controls. Mean, ± SEM, n=10). Interestingly, despite the high incidence of pericardial oedema and evidence of an impact on ventricle size and myocardial structure observed in the *hspb7* crispants, the impact on cardiovascular function in these animals was mild (**Fig. 5 and Supplementary Videos**). A small reduction in SSV in the g#1,2,3-injected animals was detected suggesting (along with blood pooling observed at 2 dpf) a small reduction in pumping efficiency (Note, however, that no impact was seen on stroke volume when ventricular dimensions were used for its calculation).

Assessment of *lmo2* crispants at 2 dpf revealed the most prominent phenotypes after injection of g#1,2,3, followed by g#2 alone (**Fig. S4**). The high mortality exhibited in the g#1,2,3-injected animals (76% in the first run), however, supported the use of g#2 for the second confirmatory run. At 4 dpf, there was a 100% incidence of heart, craniofacial, neural tube, jaw, swim bladder and yolk ball abnormalities and a high incidence of other non-cardiovascular abnormalities across all *lmo2* treatments (**Fig. 3**). The hearts of all 4 dpf *lmo2* crispants exhibited pericardial oedema, reduced heart size, lack of chamber definition and an abnormal heart shape. In addition to these cardiac specific effects, somites were poorly defined, fins small and bent, optic and otic vesicles were small and malformed, fore and midbrain boundaries were not present, brain sizes were reduced, and jaws were heavily malformed in terms of shape and size (in some individuals the presence of the jaw could not be determined). Additionally, swim bladder, liver and foregut structures could not be determined by visual inspection. The yolk ball was also excessive and the presence of yolk oedema was determined in 90% of the g#2 crispants. Histological analysis further supported the cardiac phenotype in the crispants, with observation of cardiac hypoplasia, chamber malformation and pericardial distension (**Fig. 4A and Fig. S5**). In addition, there was altered cardiomyocyte shape in the g#1,2,3, and a pyknotic nucleus structure evident in g#2-injected larvae. The severe impact of *lmo2* mutation was also reflected in significantly smaller ventricle end-diastolic and systolic dimensions, along with significantly reduced stroke volume, cardiac output and ejection fraction (**Fig.4B**). Confocal assessment of *cmlc2::DsRed2-nuc* crispants (**Fig. 4C and Fig. S6**) revealed lower cardiomyocyte numbers with frequent misshapen nuclei (estimated at 43 ±2.59 versus 79 ±2.78 in the Cas9 controls. Mean, ± SEM, n=10), along with an apparent breakdown in the uniformity of cell and DsRed2 distribution across the myocardium. The clear structural impact on the heart of *lmo2* crispants was reflected in the functional assessment (**Fig. 5 and Supplementary Videos**), with both crispant groups showing significantly reduced cardiovascular functionality compared with the Cas9 injection controls. It was also notable that the *lmo2* crispants lacked visible circulating erythrocytes, which is consistent with the key role of this gene in haematopoiesis and which also meant that blood flow and associated cardiovascular parameters were effectively zero.

## Discussion

Using human WES in conjunction with bioinformatics, three genes were identified as having a plausible link to human cardiovascular disease prevalence within large scale clinical datasets, and prioritised for further study. CRISPR/Cas9-mediated *in vivo* mutagenesis followed by morphological and functional phenotyping in zebrafish crispants was then used to reveal the role of these genes in the development and pathophysiology of the cardiovascular system. Our approach of CRISPR/Cas9-mediated multi-locus mutation in zebrafish crispants resulted in the effective mutation of all 4 genes that were targeted. Adopting multi-locus strategies to induce target gene mutation, as used here, rapidly achieve high proportions of null alleles in F0 knockouts. Such approaches, however, can also increase off-target mutations, increase double strand breaks and are not suitable for inducing site specific mutations, where targeted knockins and the creation of stable genetically modified lines may be more appropriate. Despite this, such strategies are highly beneficial for rapidly generating high-efficiency gene mutation (Kroll et al., 2021). This advantage is amplified when coupled with high throughput initial target gene identification and prioritisation, creating a potentially powerful approach for providing *in vivo* gene function data to support candidate selection as part of strategy for accelerating early drug discovery.

Supporting our approach, the positive control *gata5* crispants exhibited an expected severe and consistent impact on cardiovascular development and function at 4 dpf. Our data are consistent with the known link between *GATA5* variants and multiple human cardiovascular pathologies including familial dilated cardiomyopathy (Zhang et al., 2015) and congenital ventricular-septal defects (Wei et al., 2013). These data are also in line with previous work in zebrafish including demonstration of prominent defects in myocardial differentiation and the formation of ectopic beating myocardial tissue after loss and gain of function, respectively (Reiter et al., 1999). Here, the impact of *gata5* mutation was also evident beyond the cardiovascular system impacting various structures including the somites, fins, eyes and brain. This extracardiac impact is supported by the spatiotemporal expression of g*ata5* in developing mice (Chen et al., 2009), and by previous work in zebrafish demonstrating the central role of *gata5* in endodermal morphogenesis more broadly (Reiter et al., 2001).

The first gene target was *API5*, which encodes human apoptosis inhibitor-5 protein (Bong et al., 2020). Published evidence for a role for *API5* in human cardiovascular disease is limited to reports of a potential involvement in vascular endothelial cell apoptosis (Lu et al., 2016; Mao et al., 2020). Our *in silico* data also suggested a relatively strong association with myocardial infarction and various cardiomyopathies. *In vivo* mutation of zebrafish *api5* resulted in mild pericardial oedema and evidence of an increase in ventricle size. The latter is of particular interest given the observation of an association between *api5* downregulation and hypertrophic and dilated cardiomyopathies from the *in silico* data analysis. In the case of the former, histological analysis did not support myocardial thickening in 4 dpf zebrafish, although the absence of any clear change in cardiomyocycte number in *cmlc2::DsRed2-nuc api5*-crispants suggested that the change in ventricle size may be driven more by cardiac hypertrophy, rather than hyperplasia. Analysis of the cardiovascular phenotype in older animals would help to further clarify the mechanism(s) at play. Although published data on the function of *api5* in zebrafish are limited, it is modestly upregulated in adult zebrafish hearts following hypoxic insult (Marques et al., 2008) perhaps supporting a cardio-protective role against tissue injury. As our data suggest that *api5* does not play a critical role in early cardiovascular development, this may further support a role for *api5* in organ-system protection under conditions of physiological stress, or as a consequence of tissue injury.

*HSPB*7 encodes small heat shock protein 7, and although highly expressed in the developing and adult mammalian heart, its cardiac function remains obscure (Mercer et al., 2018). *HSPB*7 gene variants have been implicated in a range of human cardiovascular diseases including heart failure (Cappola et al., 2010; Aung et al., 2019) and dilated cardiomyopathy (Villard et al., 2011; Esslinger et al., 2017). Our *in silico* analysis supported an association of *HSPB7* with various human cardiovascular pathologies, most notably downregulation associated with various cardiomyopathies, heart failure and atrial fibrillation, and upregulation associated with dilated cardiomyopathy. A central involvement in cardiovascular development and disease is certainly supported by knockout studies in mice. Wu et al., (Wu et al., 2017) found that *Hspb7* played a critical role in development, with knockout proving embryo-lethal by around stage E12.5. Further studies on cardiac pathology in embryonic mice revealed that mutants exhibited smaller left ventricles, cardinal vein enlargement and the presence of abnormal actin bundles. Liao *et al*. (Liao et al., 2017) used an inducible-conditional knockout approach to overcome the embryo-lethal effect of *Hspb7* knockout in adult mice. These authors further revealed a critical non-developmental cardiac functional role of *Hspb7* reporting disrupted myofibrillar organisation and cardiomyocyte membrane integrity, combined with abnormal cardiac conductivity, heart arrhythmia and sudden death, likely due to a reported disruption of intercalated disc structure. Here, 4 dpf *hspb7* crispant zebrafish exhibited widespread morphological abnormalities, which included a 20% occurrence of malformed or small hearts. Furthermore, a reduction in ventricle size was also detected along with some evidence of disorganised cardiomyocyte distribution, which was reminiscent of the results previously reported in *Hspb7* knockout mice. Our data are also in broad agreement with a previous study into the role of *hspb7* in zebrafish cardiovascular development using morpholino-mediated knockdown (Rosenfeld et al., 2013). These authors reported that loss of *hspb7* function resulted in disrupted heart tube looping and ventricular cardiomyocyte development. The latter effect, specifically seen in the ventricle, was reported to be driven by reduced cardiomyocyte size, rather than number and no abnormalities of cellular ultrastructure were observed. In a more recent study, Mercer *et al*. (Mercer et al., 2018) used TALENs to generate a frameshift near the N-terminus of zebrafish *hspb7* and reported normal cardiovascular development, including timely cardiac jogging and looping. This is in contrast to the Morpholino-based knockdown work in zebrafish, previous studies in knockout mice, and the data supporting a role for *hspb7* in cardiac development generated here. In this respect, it should be noted that a crispant knockout approach generates a diversity of null alleles, which can be a drawback in disease modelling where a precise mutation needs to be duplicated. However, as our experimental objective was to reveal the consequences of the absence of the target protein, this approach can be advantageous in avoiding genetic compensation mechanisms, which can be observed in stable zebrafish knockout lines (El-Brolosy et al., 2019). Indeed, some aspects of the reported TALEN-based frame shift mutation point towards possible gene compensation, as the expression of *hsp5b* was strongly upregulated (Mercer et al., 2018). Interestingly, despite the lack of a clear morphological phenotype in the *hspb7* mutants, these animals exhibited a reduced capacity for exercise-induced cardiovascular stress, and histopathological analysis suggested an underlying pathology manifested as cardiomegaly and mild multifocal cardiac fibrosis (Mercer et al., 2018). This further suggests that a greater functional impact of *hspb7* may be observable under conditions of cardiovascular stress, something that could be tested through the use of an inducible-conditional knockout zebrafish and forced swimming assessment in adult animals.

*LMO*2 is highly conserved amongst vertebrate lineages, and encodes the Lim-domain only 2 nuclear transcriptional co-regulator crucial in early embryonic erythropoiesis and angiogenic remodelling (Chambers and Rabbitts, 2015). Beyond highlighting its well-established role in haematopoiesis and vascular development (Meng et al., 2016), our *in silico* data supported an association between *LMO2* variants and cardiomegaly and cardiomyopathy, although comparative expression data were less conclusive. CRISPR-mediated knockout in 4 dpf zebrafish here resulted in an expected absence of circulating erythrocytes, as well as widespread morphological abnormalities affecting multiple body structures. Amongst these, *lmo2* crispants uniformly exhibited reduced heart sizes, a lack of chamber definition and an abnormal heart shape. The abnormal cardiac phenotype was reinforced by clear reductions in ventricular dimensions, reduced cardiomyocyte number and disorganisation of cell distribution through the myocardium, as well significant impacts on multiple cardiovascular functional parameters. Although the role of *LMO2* in erythropoiesis is well documented, published data on cardiac-specific impacts of *LMO2* loss of function are rather more limited. Deletion of *Lmo2* in mice is embryo-lethal, with the complete failure of yolk sac erythropoiesis leading to death by around stage E10.5 (Nam and Rabbitts, 2006). Compared to wild type littermates, *Lmo2* knockout mice were reported to show no evidence of circulating erythroid cells, a small yolk sac, progressive pericardial oedema, growth retardation, significantly shortened anteroposterior axis and fewer somite pairs. Furthermore, although developing major organs were described as smaller, neurulation appeared normal, cardiac contraction was noted and there was no obvious impact on cardiac morphology (Warren et al., 1994). Knockout of *lmo2* has also previously been reported in zebrafish and appeared to result in a less severe phenotype than that observed in mice. Matrone et al. (Matrone et al., 2021) reported CRISPR/Cas9-mediated *lmo2* mutation that resulted in a 4-nucleotide insertion in exon 2 and a downstream stop codon. The resultant phenotype was described as showing mild body bending, reduced skin pigmentation, fewer circulating red blood cells, mild pericardial oedema, and mild cephalomegaly at 72hpf. Weiss *et al*. (Weiss et al., 2012) also reported severe head oedema and impaired optic fissure closure in 2 dpf *lmo2* mutants isolated from an ENU-mutagenesis screen. The reason for this apparent difference in severity versus our *lmo2* crispants is not clear, however, the complete absence of visible erythrocytes in 29/30 crispants assessed for cardiovascular function (1 animal had around 10 visible erythrocytes in the dorsal aorta) does support the extremely effective knockout out of *lmo2* function. Moreover, in our g#2 injected animals, an absence of blood flow was apparent in all resultant crispants. Additionally, 73% of these animals showed pericardial oedema, 82% misshapen hearts, and 11% a lack of heart chamber definition at 2 dpf suggesting the observed cardiac phenotype was initiated at a relatively early stage of development. Although it is unknown if *lmo2* is expressed in cardiomyocytes or endocardial cells themselves, it is well known that fluid forces can profoundly affect cardiac structural development. Reviewed by Sidhwani and Yelon (Sidhwani and Yelon, 2019), blood flow has, for example, been shown to impact endothelial cell number and polarity, cardiac chamber morphology, cardiomyocyte shape, size and myofibril maturation, as well as endocardial cell development and morphology, atrioventricular valve formation and ventricular trabeculation. Consequently the absence of circulating erythrocytes in our *lmo2* cripsants could have significantly affected the structure and function of the developing heart. At this point, we also cannot exclude the possibility of off-target effects contributing to the observed phenotype at 4dpf, particularly in the g#1,2,3 crispants. Although off-target effects are potentially more likely in this group due to the targeting of 3 loci on *lmo2*, the demonstration of a consistent phenotype across 3 separate gRNA injection groups at 2 dpf, and between the g#1,2,3 and g#2 injected animals at 4 dpf does support the notion that *lmo2* gene function was specifically impaired. Regarding the wider developmental impact of *lmo2* deletion in our crispants, although data in zebrafish are limited, it has been reported that in embryonic mice there is consistent expression of *Lmo2* in multiple non-haematopoietic/vascular tissues in early-mid gestation including brain, eyes, somites, liver, limb buds, tail buds and developing limbs. Furthermore, the organisation of expression in an anterior/posterior pattern was interpreted as a likely involvement in some major patterning activities during early development (Calero-Nieto et al., 2013), perhaps hinting at a wider developmental role for this gene. As suggested with *hspb7*, this is a case where the generation of an inducible-conditional knockout model may help to delineate the role of *lmo2* outside that of early development.

In conclusion, using a novel combination of *in silico* analysis of clinical data with *in vivo* assessment of CRISPR/Cas9-mediated mutation in zebrafish crispants we have generated data to strongly support the role of *api5, hspb7* and *lmo2* (and *gata5*) in vertebrate cardiovascular development and function, and by inference, the potential effect of loss of gene function on subsequent organ system pathophysiology. The approach we used allowed rapid screening of the impact of target gene mutation on embryo-larval development and organ system function, thus facilitating early drug discovery by providing *in vivo* data to support the selection and prioritisation of candidate genes for further investigation. Although this approach provides *in vivo* data on gene function within just a few days, the assessment of phenotypes in early zebrafish larvae has some limitations. For example, delineation of the developmental and adult roles of target genes is more difficult, and embryo-lethal phenotypes would preclude the assessment of gene function at later life stages. In this respect, the generation of conditional inducible knockouts in older zebrafish would be a highly valuable next step as this would provide data on gene function in a more directly translatable model of adult human heart failure. Moreover, assessing organ system functionality at rest may also be relatively insensitive, whereas the use of experimental paradigms, in which cardiovascular stress is applied, may prove more revealing when relating the loss of gene function to real clinical outcomes.

## Acknowledgements

The authors would like to thank the staff in the Aquatic Resources Centre at the University of Exeter, for the supply and maintenance of the zebrafish. In addition, the *cmlc2::DsRed2-nuc* fish were generously provided by Dr. Geoff Burns, Boston Children’s Hospital and Harvard Medical School. The authors would also like to thank Drs Keith Cheng and Alex Yu-Shun Lin, at Pennsylvania State University College of Medicine, for their advice on the histological analysis.

## Competing interests

S.R., M.B-Y., A.W. and E.M. are employees of AstraZeneca PLC. There are no other potential conflicts of interest.

## Funding

This work was supported by AstraZeneca, a Royal Society short Industry Fellowship (SIF\R2\192004) awarded to M.J.W. and the University of Exeter College of Life and Environmental Sciences. In addition, research in the Scholpp lab is supported by the BBSRC (Research Grant, BB/S016295/1) and by the Living Systems Institute, University of Exeter.

## Data availability

All data associated with this study are available in the article or the supplementary materials.

## Author contributions statement

S.R., M.B-Y. and M.J.H. conceived the project and developed the concept. S.R., M.B-Y., M.J.H, S.S., C.R.T. and M.J.W. obtained the funding. A.W., E.M. and M.B. undertook the *in silico* work. M.J.W., Y.O., J.S.B., M.J.H., S.S., C.R.T. and M.B-Y. designed the zebrafish experiments. M.J.W., Y.O., J.S.B. and A.T. undertook the zebrafish experiments and analysed the resultant data. Data interpretation and manuscript production was undertaken by all authors.

## List of supplementary information

**Table S1**. Summary of literature data on the structure and function of the target genes identified for zebrafish-based *in vivo* investigation in the current study

**Table S2**. List of morphological endpoints scored as a simple yes/no at 2 dpf to facilitate selection of treatment groups for more in depth morphological scoring and functional phenotype analysis at 4 dpf

**Table S3**. List of morphological endpoints scored in severity from 1 (severe) to 5 (normal) at 4 dpf Scoring criteria adapted from those of Gustafson *et al*., (2012) and Ball *et al*., (2014)

**Table S4**. The sequence of fixation and embedding steps applied to 4 dpf zebrafish using an automated tissue embedder

**Table S5**. The sequence of H&E staining steps applied to 4 dpf zebrafish using an automated stainer

**Fig. S1**. gRNA design for each of the 4 targeted genes

**Fig. S2**. GeneMANIA network analysis using API5, HSPB7, LMO2 and GATA5 as input genes

**Fig. S3**. Transcriptomics data analysis using *API5, HSPB7, LMO2* and *GATA5* as input genes

**Fig. S4**. Comparison of morphological endpoints measured in 2 dpf animals

**Figure S5**. Haematoxylin and eosin stained coronal sections through the hearts of 4 dpf zebrafish crispants

**Fig. S6**. Confocal maximum intensity projection images of hearts from *cmlc2::DsRed2-nuc* larvae in which the targets genes were mutated

**Supplementary videos**. Videos of cardiovascular function to accompany **Figure 5** in the main manuscript

**Supplementary datasets. Data S1**. Data S1_Network Analysis.xlsx; **Data S2**. Data S2_Differential Expression.xlsx; **Data S3**. Data S3_Genetic Associations.xlsx; **Data S4**. Data S4_Homologies.xlsx

## Notes

### Competing Interest Statement

SR, MB, AW and EM are employees of AstraZeneca PLC. There are no other potential conflicts of interest.

### Summary of Updates

Addition of more data and redrafting of manuscript.

