## Supplementary information for "A combined human *in silico* and CRISPR/Cas9-mediated *in vivo* zebrafish based approach for supporting gene target validation in early drug discovery"

### SUPPLEMENTARY TABLES

**Table S1.** Summary of literature data on the structure and function of the target genes identified for zebrafish-based *in vivo* investigation in the current study

| OFFICIAL GENE SYMBOL (ALIASES) | GENE NAME(S) | PROTEIN STRUCTURE & FUNCTION(S) | BIOLOGICAL & PHYSIOLOGICAL FUNCTION(S) |
|---|---|---|---|
| ***API5***<br><br>***(AAC-11; FIF; MIG8)*** | Apoptosis Inhibitor 5; Anti-apoptosis clone 11, Fibroblast Growth Factor 2-Interacting Factor;<br>Cell migration-inducing gene 8 | • Belongs to the Inhibitor of apoptosis (IAP) family characterized originally as physical inhibitors of caspases (Salvesen et al 2002)<br>• May act as a scaffold for multiprotein complexes. Contains a multiple helices, forming HEAT (Huntingtin, elongation factor 3, PR65/A, TOR)-like and ARM (Armadillo)-like repeats that mediate protein–protein interactions (Han et al 2012),<br>• The C-terminal is structurally similar to the core region of the U-box-containing ubiquitin ligase E4 protein Ufd2p (Han et al 2012) | <u>*Apoptosis*</u><br>• Prevents apoptosis after growth factor deprivation (Tewari et al 1997)<br>• Prevents cell death by negatively regulating E2F1 transcription factor-induced apoptosis (Morris et al 2006). Contributes to E2F1 control of the G1/S cell cycle phase transition via facilitating E2F1 recruitment onto its target promoters and thus E2F1 target gene transcription (Garcia-Jove Navarro et al 2013)<br>• Regulates Acinus (ACIN1), a protein involved in chromatin condensation and DNA fragmentation during apoptosis protecting it from caspase 3 cleavage (Rigou et al 2009)<br>• Binds to and inhibits caspase-2, a key regulator of cell death, autophagy, genomic stability and ageing (Imre et al 2017)<br>• Interacts with FGF2 (Van den Berghe et al 2000) and upregulates FGF2 signaling through a FGFR1/PKCdelta/ERK effector pathway triggering degradation of the proapoptotic molecule BIM (Noh et al 2014)<br>• Regulates miR-1 induced apoptosis (Li et al 2015)<br>• Phosphorylated by PIM2 kinase and inhibits apoptosis in hepatocellular carcinoma cells through NF-κB (nuclear factor-κB) (Ren et al 2010)<br>• Downregulated in cardiomyocytes undergoing $H_2O_2$-induced apoptosis (Wan et al 2016)<br><u>*Tumorigenesis*</u><br>• Upregulated in human carcinomas in vivo (Koci et al 2012)<br>• Inhibition increases anti-cancer drug sensitivity in various cancer cells (Rigou et al 2009) |

| | | | |
|---|---|---|---|
| | | | • Increases the metastatic capacity of tumor cells by upregulating MMP levels via activation of the Erk signaling pathway (Song et al 2015)<br>• Acts as an immune escape gene in tumors by rendering them resistant to apoptosis triggered by tumor antigen-specific T cells (Noh et al 2014)<br>*Immune response*<br>• Acts as a danger-associated molecular pattern (DAMP) and stimulate the activation of immune response. Mediates TLR4-dependent activation of antigen presenting cells (Kim et al 2017) |
| ***HSPB7***<br><br>***(CVHSP)*** | Heat Shock Protein Family B (Small) Member 7; Cardiovascular Heat Shock Protein | • Member of the small heat shock proteins (HSPB) family of molecular chaperones. HSPBs typically associate early with misfolded proteins (Mymrikov et al 2011)<br>• Prevents aggregation of proteins with expanded polyglutamine (polyQ) stretches (Vos et al 2010)<br>• Binds large sarcomeric proteins Filamin C and Titin in cardiac cells (Mercer et al 2018) | *Cardiac function*<br>• Selectively expressed in cardiac and skeletal muscle<br>• Common genetic variants associate with human HF (Cappola et al 2010)<br>• Modulates actin filament assembly in heart (Wu et al 2017)<br>• Cardiac-specific inducible HSPB7 KO mice display cardiac arrhythmia with abnormal conduction, rapid HF and sudden death accompanied by a downregulation of intercalated disc proteins and enrichment in Filamin C protein aggregates (Liao et al 2017)<br>• Global depletion of Hspb7 in zebrafish disrupts normal cardiac morphogenesis (Rosenfeld et al 2013)<br>• Loss of HSPB7 in zebrafish or human cardiomyocytes leads to enhanced autophagic pathways (Mercer et al 2018)<br>• Dramatically increased in the heart and blood plasma immediately after myocardial infarction (Rüdebusch et al 2017)<br>*Tumorigenesis*<br>• May act as a tumor suppressor in the p53 pathway (Lin et al 2014) |
| ***LMO2***<br><br>***(TTG2; RBTN2; RHOM2)*** | LIM Domain Only 2; Rhombotin-2 | • Belongs to the LIM-Only group of LIM domain protein superfamily.<br>• The LIM-fingers are distinct from the zinc-fingers of transcription factors such as *GATA*, as LIM-domain-only proteins do not directly bind to DNA (8)<br>• Mediates transactivation via protein-protein interactions with other transcriptional factors. | *Angiogenesis*<br>• Regulates endothelial proliferation and angiogenesis *in vitro* and is required for angiogenesis and tissue healing *in vivo* (Meng et al 2016)<br>• Promotes endothelial cell migration via modulation of Sphk1 (Sphingosine Kinase). *Lmo2* KD reduced Lmo2-Sphk1 gene interaction, impaired intersegmental vessels formation, and reduced cell migration ( Matrone et al 2017)<br>• Angiopoietin-2 and VE-cadherin, both major regulators of angiogenesis, are direct transcriptional targets of LMO2-complexes with bHLH transcription factors TAL1 and/or LYL1 in endothelial cells (Deleuze et |

| | | | |
|---|---|---|---|
| | | | al. 2007, 2012)<br>• Regulate VEGF-induced angiogenesis and lymphangiogenesis via NRP2 (Coma et al 2013)<br><u>*Hematopoiesis/Erythropoiesis*</u><br>• Homozygous inactivation in mice led to the complete absence of yolk sac erythropoiesis and early embryonic lethality (Warren et al. 1994)<br>• Acts with TAL1/SCL to regulate erythropoiesis (Valge-Archer et al 1994)<br><u>*Tumorigenesis*</u><br>• Attenuates tumor growth by targeting the Wnt signaling pathway in breast and colorectal cancer (Liu et al 2016)<br>• Regulator of neo-vascularization of tumors (Yamada et al. 2002)<br>• Acts as a proto-oncogene in T cells (Mao et al 1997). Recurrent findings of interstitial deletions and translocations involving *LMO2* in T cell leukaemias. |

**References from Table S1**

***API5:***

Wan C, Xiang J, Li Y, Guo D. Differential Gene Expression Patterns in Chicken Cardiomyocytes during Hydrogen Peroxide-Induced Apoptosis. Plos one. 2016;11(1):e0147950. DOI: 10.1371/journal.pone.0147950.

Song KH, Kim SH, Noh KH, et al. Apoptosis Inhibitor 5 Increases Metastasis via Erk-mediated MMP expression. BMB Reports. 2015 Jun;48(6):330-335. DOI: 10.5483/bmbrep.2015.48.6.139.

Li D, Liu Y, Li H, et al. MicroRNA-1 promotes apoptosis of hepatocarcinoma cells by targeting apoptosis inhibitor-5 (API-5). FEBS Letters. 2015 Jan;589(1):68-76. DOI: 10.1016/j.febslet.2014.11.025.

Noh KH, Kim SH, Kim JH, et al. API5 confers tumoral immune escape through FGF2-dependent cell survival pathway. Cancer Research. 2014 Jul;74(13):3556-3566. DOI: 10.1158/0008-5472.can-13-3225.

Garcia-Jove Navarro M, Basset C, Arcondéguy T, et al. Api5 contributes to E2F1 control of the G1/S cell cycle phase transition. Plos one. 2013 ;8(8):e71443. DOI: 10.1371/journal.pone.0071443.

Koci L, Chlebova K, Hyzdalova M, et al. Apoptosis inhibitor 5 (API-5; AAC-11; FIF) is upregulated in human carcinomas in vivo. Oncology Letters. 2012 Apr;3(4):913-916. DOI: 10.3892/ol.2012.593.

Han BG, Kim KH, Lee SJ, et al. Helical repeat structure of apoptosis inhibitor 5 reveals protein-protein interaction modules. The Journal of Biological Chemistry. 2012 Mar;287(14):10727-10737. DOI: 10.1074/jbc.m111.317594.

Morris EJ, Michaud WA, Ji JY, et al. Functional identification of Api5 as a suppressor of E2F-dependent apoptosis in vivo. Plos Genetics. 2006 Nov;2(11):e196. DOI: 10.1371/journal.pgen.0020196.

Tewari M, Yu M, Ross B, et al. AAC-11, a novel cDNA that inhibits apoptosis after growth factor withdrawal. Cancer Research. 1997 Sep;57(18):4063-4069.

Ren K, Zhang W, Shi Y, Gong J. Pim-2 activates API-5 to inhibit the apoptosis of hepatocellular carcinoma cells through NF-kappaB pathway. Pathology Oncology Research : POR. 2010 Jun;16(2):229-237. DOI: 10.1007/s12253-009-9215-4.

Salvesen GS, Duckett CS. IAP proteins: blocking the road to death's door. Nature reviews. Molecular Cell Biology. 2002 Jun;3(6):401-410. DOI: 10.1038/nrm830.

Van den Berghe L, Laurell H, Huez I, et al. FIF [fibroblast growth factor-2 (FGF-2)-interacting-factor], a nuclear putatively antiapoptotic factor, interacts specifically with FGF-2. Molecular Endocrinology (Baltimore, Md.). 2000 Nov;14(11):1709-1724. DOI: 10.1210/mend.14.11.0556.

Rigou P, Piddubnyak V, Faye A, et al. The antiapoptotic protein AAC-11 interacts with and regulates Acinus-mediated DNA fragmentation. The EMBO Journal. 2009 Jun;28(11):1576-1588. DOI: 10.1038/emboj.2009.106.

Kim YS, Park HJ, Park JH, et al. A novel function of API5 (apoptosis inhibitor 5), TLR4-dependent activation of antigen presenting cells. Oncoimmunology. 2018 ;7(10):e1472187. DOI: 10.1080/2162402x.2018.1472187.

Imre G, Berthelet J, Heering J, et al. Apoptosis inhibitor 5 is an endogenous inhibitor of caspase-2. EMBO Reports. 2017 May;18(5):733-744. DOI: 10.15252/embr.201643744.

***HSPB7:***

Liao WC, Juo LY, Shih YL, Chen YH, Yan YT. HSPB7 prevents cardiac conduction system defect through maintaining intercalated disc integrity. PLoS Genet. 2017 Aug 21;13(8):e1006984. doi: 10.1371/journal.pgen.1006984. PMID: 28827800; PMCID: PMC5587339.

Mymrikov EV, Seit-Nebi AS, Gusev NB. Large potentials of small heat shock proteins. Physiol Rev. 2011 Oct;91(4):1123-59. doi: 10.1152/physrev.00023.2010. PMID: 22013208.

Wu T, Mu Y, Bogomolovas J, Fang X, Veevers J, Nowak RB, Pappas CT, Gregorio CC, Evans SM, Fowler VM, Chen J. HSPB7 is indispensable for heart development by modulating actin filament assembly. Proc Natl Acad Sci U S A. 2017 Nov 7;114(45):11956-11961. doi: 10.1073/pnas.1713763114. Epub 2017 Oct 23. PMID: 29078393; PMCID: PMC5692592.

Cappola TP, Li M, He J, et al. Common variants in HSPB7 and FRMD4B associated with advanced heart failure. Circulation. Cardiovascular Genetics. 2010 Apr;3(2):147-154. DOI: 10.1161/circgenetics.109.898395.

Rüdebusch J, Benkner A, Poesch A, et al. Dynamic adaptation of myocardial proteome during heart failure development. Plos one. 2017 ;12(10):e0185915. DOI: 10.1371/journal.pone.0185915.

Vos MJ, Zijlstra MP, Kanon B, van Waarde-Verhagen MA, Brunt ER, Oosterveld-Hut HM, Carra S, Sibon OC, Kampinga HH. HSPB7 is the most potent polyQ aggregation suppressor within the HSPB family of molecular chaperones. Hum Mol Genet. 2010 Dec 1;19(23):4677-93. doi: 10.1093/hmg/ddq398. Epub 2010 Sep 15. PMID: 20843828.

Rosenfeld GE, Mercer EJ, Mason CE, Evans T. Small heat shock proteins Hspb7 and Hspb12 regulate early steps of cardiac morphogenesis. Developmental Biology. 2013 Sep;381(2):389-400. DOI: 10.1016/j.ydbio.2013.06.025.

Mercer EJ, Lin YF, Cohen-Gould L, Evans T. Hspb7 is a cardioprotective chaperone facilitating sarcomeric proteostasis. Developmental Biology. 2018 Mar;435(1):41-55. DOI: 10.1016/j.ydbio.2018.01.005.

Lin J, Deng Z, Tanikawa C, Shuin T, Miki T, Matsuda K, Nakamura Y. Downregulation of the tumor suppressor HSPB7, involved in the p53 pathway, in renal cell carcinoma by hypermethylation. Int J Oncol. 2014 May;44(5):1490-8. doi: 10.3892/ijo.2014.2314. Epub 2014 Feb 27. PMID: 24585183; PMCID: PMC4027944.

***LMO2:***

Chambers J, Rabbitts TH. LMO2 at 25 years: a paradigm of chromosomal translocation proteins. Open Biol. 2015 Jun;5(6):150062. doi: 10.1098/rsob.150062. PMID: 26108219; PMCID: PMC4632508.

Matrone G, Meng S, Gu Q, et al. Lmo2 (LIM-Domain-Only 2) Modulates Sphk1 (Sphingosine Kinase) and Promotes Endothelial Cell Migration. Arteriosclerosis, Thrombosis, and Vascular Biology. 2017 Oct;37(10):1860-1868. DOI: 10.1161/atvbaha.117.309609.
Deleuze V, El-Hajj R, Chalhoub E, Dohet C, Pinet V, Couttet P, Mathieu D. Angiopoietin-2 is a direct transcriptional target of TAL1, LYL1 and LMO2 in endothelial cells. PLoS One. 2012;7(7):e40484. doi: 10.1371/journal.pone.0040484. Epub 2012 Jul 6. PMID: 22792348; PMCID: PMC3391236.
Deleuze V, Chalhoub E, El-Hajj R, Dohet C, Le Clech M, Couraud PO, Huber P, Mathieu D. TAL-1/SCL and its partners E47 and LMO2 up-regulate VE-cadherin expression in endothelial cells. Mol Cell Biol. 2007 Apr;27(7):2687-97. doi: 10.1128/MCB.00493-06. Epub 2007 Jan 22. PMID: 17242194; PMCID: PMC1899886.
Coma S, Allard-Ratick M, Akino T, van Meeteren LA, Mammoto A, Klagsbrun M. GATA2 and Lmo2 control angiogenesis and lymphangiogenesis via direct transcriptional regulation of neuropilin-2. Angiogenesis. 2013 Oct;16(4):939-52. doi: 10.1007/s10456-013-9370-9. Epub 2013 Jul 28. PMID: 23892628; PMCID: PMC3793839.
Liu Y, Huang D, Wang Z, Wu C, Zhang Z, Wang D, Li Z, Zhu T, Yang S, Sun W. LMO2 attenuates tumor growth by targeting the Wnt signaling pathway in breast and colorectal cancer. Sci Rep. 2016 Oct 25;6:36050. doi: 10.1038/srep36050. PMID: 27779255; PMCID: PMC5078767.
Warren AJ, Colledge WH, Carlton MB, Evans MJ, Smith AJ, Rabbitts TH. The oncogenic cysteine-rich LIM domain protein rbtn2 is essential for erythroid development. Cell. 1994 Jul 15;78(1):45-57. doi: 10.1016/0092-8674(94)90571-1. PMID: 8033210.
Yamada, Y., Pannell, R., Forster, A. et al. The LIM-domain protein Lmo2 is a key regulator of tumour angiogenesis: a new anti-angiogenesis drug target. Oncogene 21, 1309–1315 (2002). https://doi.org/10.1038/sj.onc.1205285
Valge-Archer VE, Osada H, Warren AJ, Forster A, Li J, Baer R, Rabbitts TH. The LIM protein RBTN2 and the basic helix-loop-helix protein TAL1 are present in a complex in erythroid cells. Proc Natl Acad Sci U S A. 1994 Aug 30;91(18):8617-21. doi: 10.1073/pnas.91.18.8617. PMID: 8078932; PMCID: PMC44657.
Meng S, Matrone G, Lv J, Chen K, Wong WT, Cooke JP. LIM Domain Only 2 Regulates Endothelial Proliferation, Angiogenesis, and Tissue Regeneration. J Am Heart Assoc. 2016 Oct 6;5(10):e004117. doi: 10.1161/JAHA.116.004117. PMID: 27792641; PMCID: PMC5121509.
Visvader JE, Mao X, Fujiwara Y, Hahm K, Orkin SH. The LIM-domain binding protein Ldb1 and its partner LMO2 act as negative regulators of erythroid differentiation. Proc Natl Acad Sci U S A. 1997 Dec 9;94(25):13707-12. doi: 10.1073/pnas.94.25.13707. PMID: 9391090; PMCID: PMC28370.
Mao S, Neale GA, Goorha RM. T-cell proto-oncogene rhombotin-2 is a complex transcription regulator containing multiple activation and repression domains. J Biol Chem. 1997 Feb 28;272(9):5594-9. doi: 10.1074/jbc.272.9.5594. PMID: 9038167.

**Table S2.** List of morphological endpoints scored as a simple yes/no at 2dpf to facilitate selection of treatment groups for more in depth morphological scoring and functional phenotype analysis at 4dpf.

| Endpoint | Score criteria | Score | Scored yes/no with notes |
|---|---|---|---|
| **Body shape** | Bent/curved | Yes/No | N/A if not dechorionated* |
| **Somites** | Poor definition | Yes/No | |
| **Notochord** | Present / absent | Yes/No | |
| **Tail** | Bent/curved | Yes/No | N/A if not dechorionated* |
| **Heart** | No heartbeat | Yes/No | |
| | Slow heartbeat | Yes/No | |
| | Swollen pericardial sac | Yes/No | |
| | Chambers not well defined | Yes/No | |
| | Misshapen | Yes/No | |
| **Face** | Optic vesicle small | Yes/No | |
| | Otic vesicles small | Yes/No | |
| | Facial oedema | Yes/No | |
| **Neural** | Irregular shape | Yes/No | Indicate fore, mid or hindbrain |
| **Jaw/Arches** | Present | Yes/No | |
| **Body Oedema** | Oedema present | Yes/No | In addition to heart or facial tissue |

*Need to be hatched or dechlorinated in advance otherwise a bent body form will persist

**Table S3.** List of morphological endpoints scored in severity from 1 (severe) to 5 (normal) at 4dpf. Scoring criteria adapted from those of Gustafson *et al*., (2012) and Ball *et al*., (2014).

| Endpoint | Score criteria | Score | Notes ~ scored 1 to 5 (normal)* |
|---|---|---|---|
| **Body shape** | Bent/curved | Yes/No | |
| | Body length | N/A | Standard length in mm |
| **Somites** | Poor definition | Yes/No | |
| | Small./ short | Yes/No | |
| | Missing | Yes/No | |
| | Cloudy | Yes/No | |
| **Notochord** | Shortened | Yes/No | |
| | Folded in Tail | Yes/No | |
| | No cellular differentiation | Yes/No | |
| | Wavy/Kinked | Yes/No | |
| | Poorly defined | Yes/No | |
| **Tail** | Kinked | Yes/No | |
| | Bent/Curved | Yes/No | |
| | Short | Yes/No | |
| **Fins** | Small | Yes/No | Indicate affected fin(s) |
| | Irregular Edge | Yes/No | |
| | Bent | Yes/No | |
| | Cloudy | Yes/No | |
| | Eroding | Yes/No | |
| **Heart** | Slow Heartbeat | Yes/No | |
| | No Heartbeat | Yes/No | |
| | Pericardial Sac-Swollen | Yes/No | |
| | Chambers not well defined | Yes/No | |
| | Misshapen | Yes/No | |
| **Face** | Optic Vesicle – Small | Yes/No | |
| | Optic Vesicle – Misshapen | Yes/No | |
| | Otic Vesicles – Small | Yes/No | |
| | Olfactory Region – Reduced | Yes/No | |
| | Olfactory Region - Not Present | Yes/No | |
| | Facial Oedema or Hypoplasia | Yes/No | |
| **Neural Tube** | Irregular Shape | Yes/No | Indicate irregular region(s) |
| | Reduced/Compressed | Yes/No | |
| **Arches/Jaw#** | Arches - Not Evident | Yes/No | |
| | Arches - Irregular Shape | Yes/No | |
| | Arches – Deficient | Yes/No | |
| | Jaw - Not Evident | Yes/No | |
| | Jaw - Deficient | Yes/No | Indicate irregular region(s) |
| | Jaw - Enlarged/ excessive | Yes/No | |
| | Jaw - Irregular Shape | Yes/No | |
| **Liver** | Enlarged | Yes/No | |
| | Not evident | Yes/No | |
| **Yolk ball** | Remnant excessive | Yes/No | |

#Scoring of the jaw is variable at 4dpf and he jaw at 4dpf, will be more variable at that stage.

**Table S4.** Sequence of fixation and embedding steps applied to 4dpf zebrafish using an automated tissue embedder.

| Solution | Duration | Temperature (ºC) | Vacuum |
|---|---|---|---|
| 70% isopropanol | 60min | 30 | on |
| 90% isopropanol | 60min | 30 | on |
| 95% isopropanol | 60min | 30 | on |
| 100% isopropanol | 60min | 30 | on |
| 100% isopropanol | 60min | 30 | on |
| 100% isopropanol | 60min | 30 | on |
| Xylene | 60min | 30 | on |
| Xylene | 60min | 30 | on |
| Xylene | 60min | 30 | on |
| Wax | 80min | 62 | on |
| Wax | 80min | 62 | on |
| Wax | 80min | 62 | on |

**Table S5.** Sequence of H&E staining steps applied to 4dpf zebrafish using an automated stainer.

| Solution | Duration |
|---|---|
| Histoclear | 5min |
| Histoclear | 5min |
| 100% IMS | 2min |
| 90% IMS | 2min |
| 80% IMS | 2min |
| Tap water (running) | 2min |
| Harris Haematoxylin non-acidified | 15min |
| Tap water (running) | 2min |
| Acid Alcohol | 5sec |
| Tap water (running) | 30sec |
| Ammoniated alcohol | 30sec |
| Tap water (running) | 30sec |
| Eosin Y Aqueous | 15sec |
| Tap water (running) | 30sec |
| 80% IMS | 30sec |
| 90% IMS | 1min |
| 95% IMS | 1min |
| 100% IMS | 2min |
| 100% Ethanol | 2min |
| Histoclear | 2min |
| Histoclear | 2min |

IMS: Industrial Methylated Spirit

### SUPPLEMENTARY FIGURES

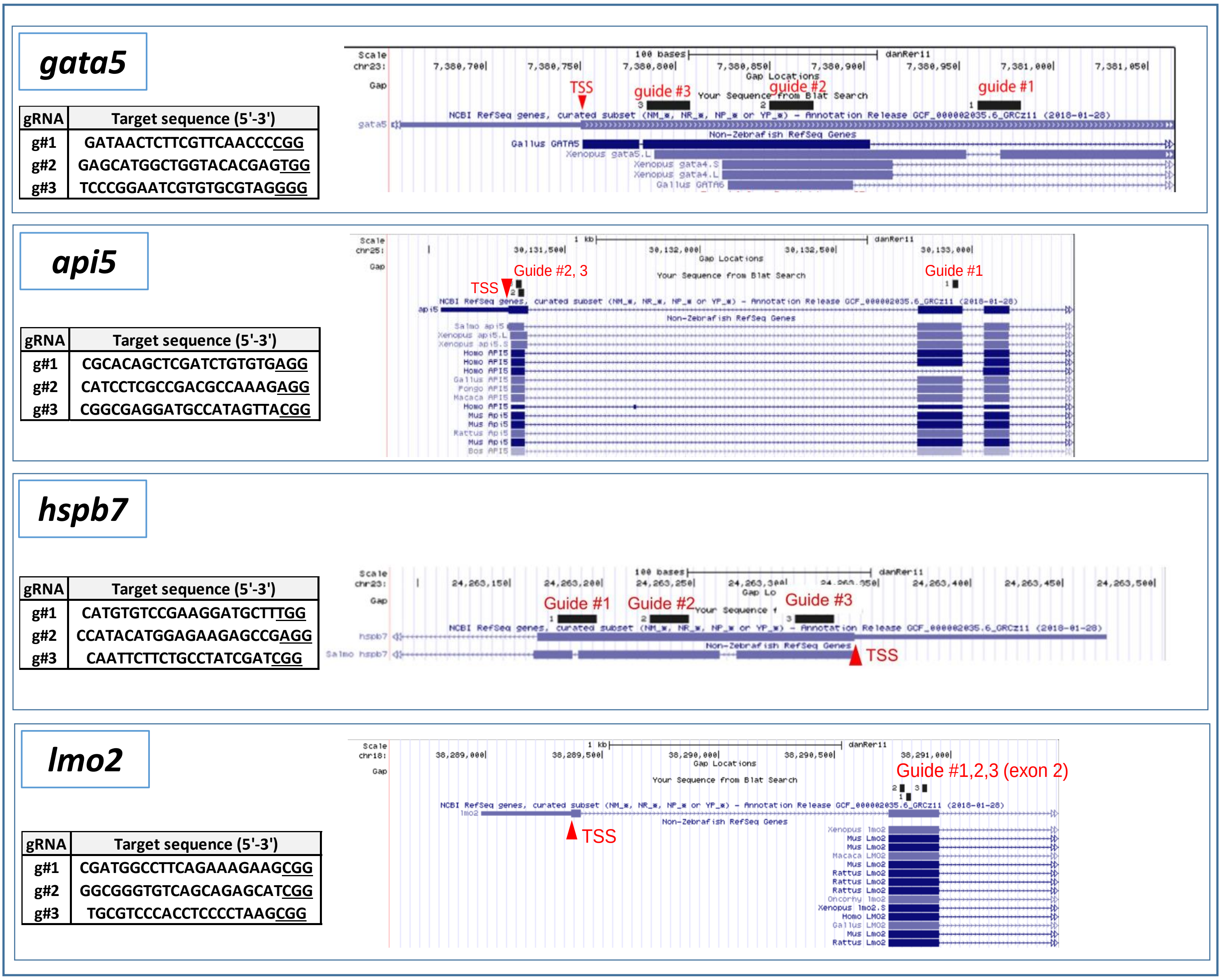


**Fig. S1. gRNA design and mutation efficient for each of the 4 targeted genes.** For each gene in turn, the left hand table shows the sequence of each designed gRNA (g#1,2,3 encompassed all three gRNAs in one injection), and the right hand image shows the position of each of these gRNA sequences within the coding exons of each target gene. The translation start site is indicated by TSS and the adjoining arrow, and other species orthologues are shown for comparison. The three guides were selected to target distinct regions of each gene to assess the varying impact, and also to ensure effective disruption in the case of the gRNA1,2,3-injected group. Original images obtained using the UCS genome browser (https://genome.ucsc.edu).

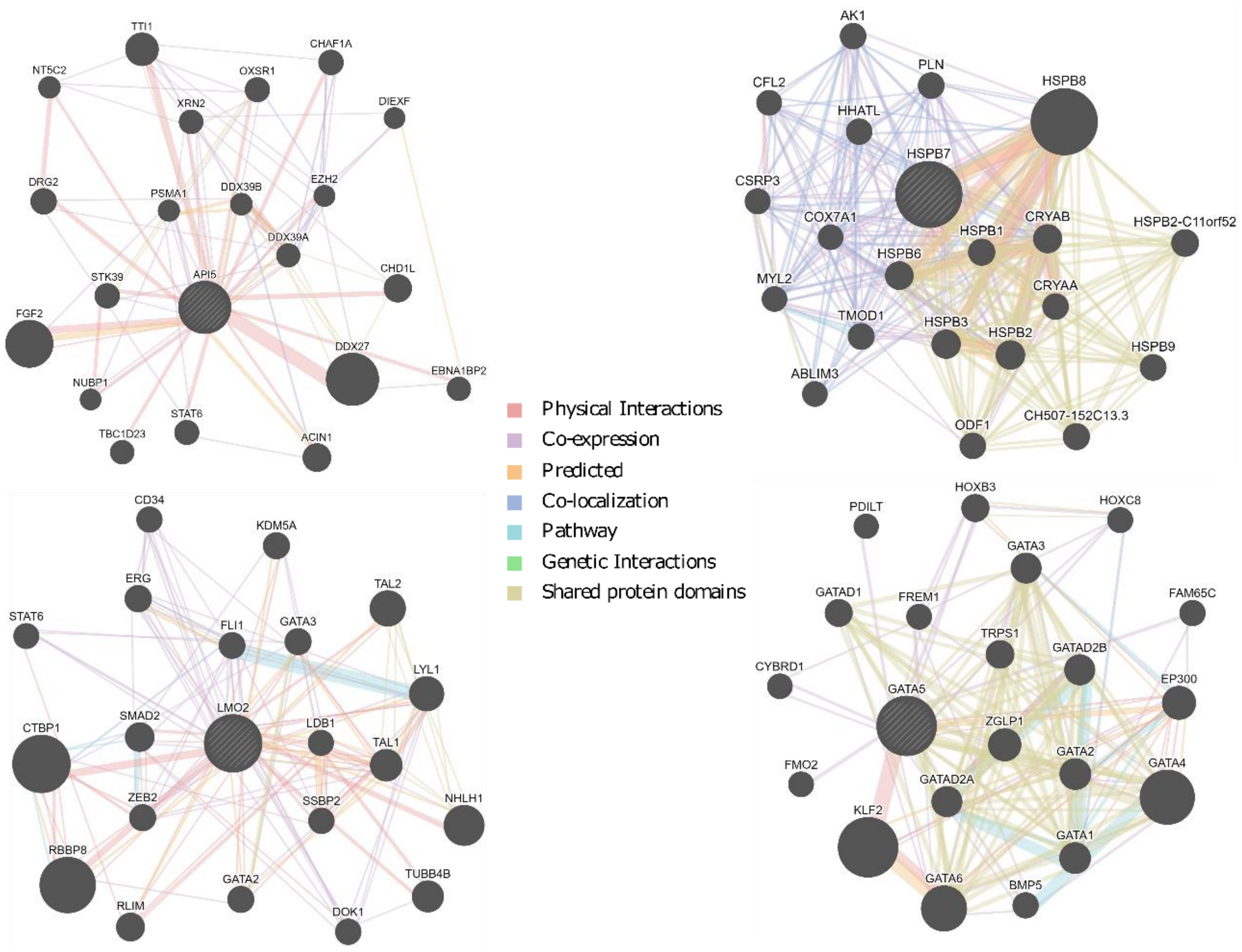


**Fig. S2. GeneMANIA network analysis using *API5*, *HSPB7*, *LMO2* and *GATA5* as input genes.** GeneMANIA uses very large set of functional association data including protein and genetic interactions, pathways, co-expression, co-localization and protein domain similarity (Warde-Farley *et al*., 2010. *Nucleic Acids Res*, 38, W214-20). Colour of network edges indicate evidence types.

### API5 – Tissue distribution in humans

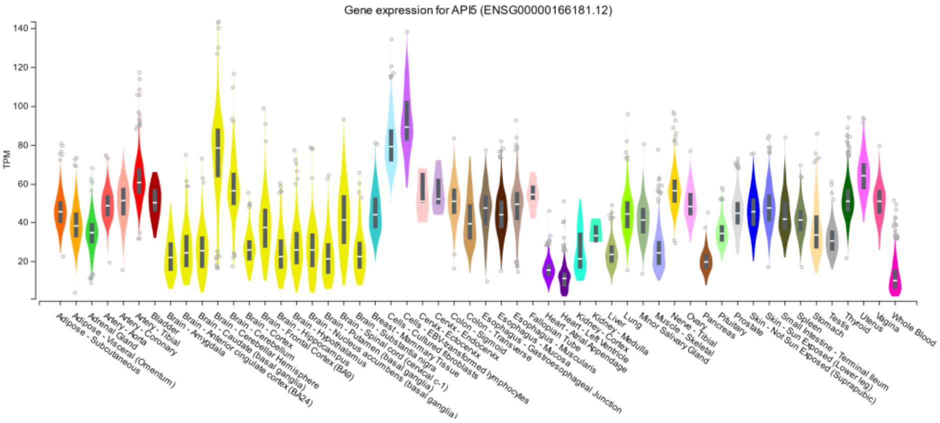


### API5 – Tissue distribution in humans

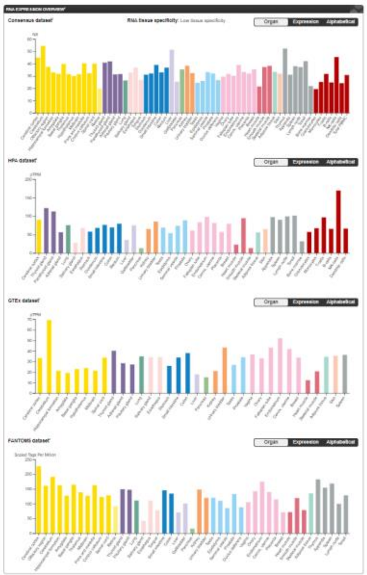

### API5 – Cell type expression in humans

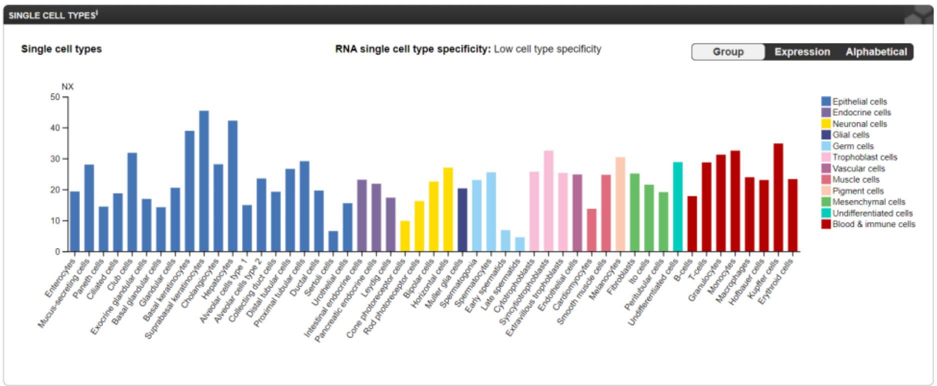


### API5 – Cellular expression in human heart

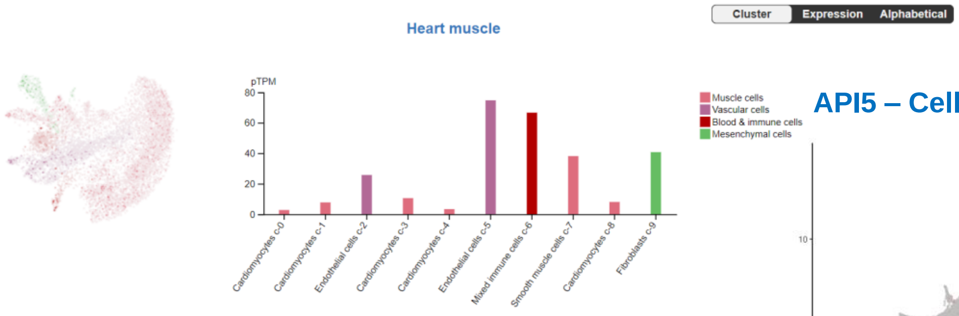


### API5 – Cellular expression in human heart

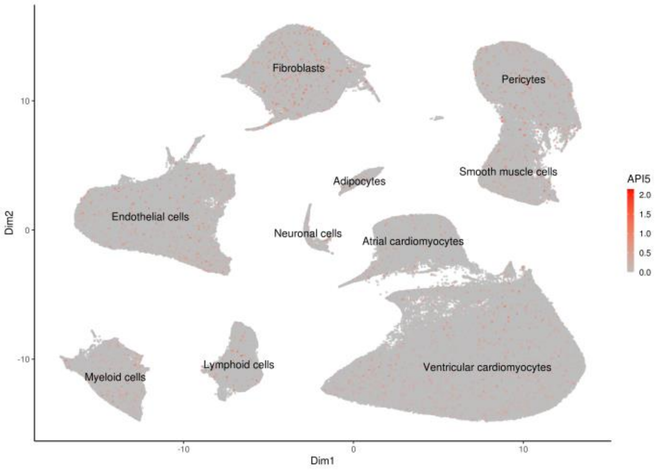


**Fig. S3A. Transcriptomics data analysis on *API5*.** Data shown are the tissue and single cell type mRNA expression levels based on RNA sequencing data sets from GTExPortal (https://gtexportal.org/home/), the Human Protein Atlas (https://www.proteinatlas.org/) and the Heart Cell Atlas (https://www.heartcellatlas.org/).

### HSPB7 – Tissue distribution in humans

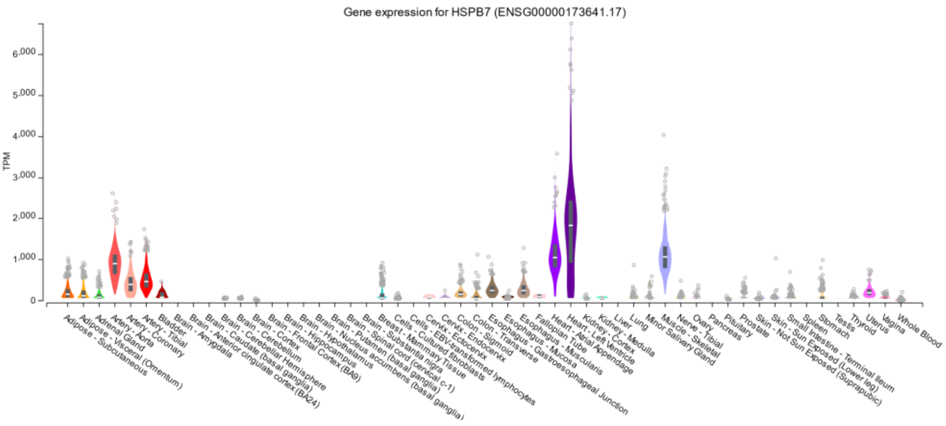


### HSPB7 – Tissue distribution in humans

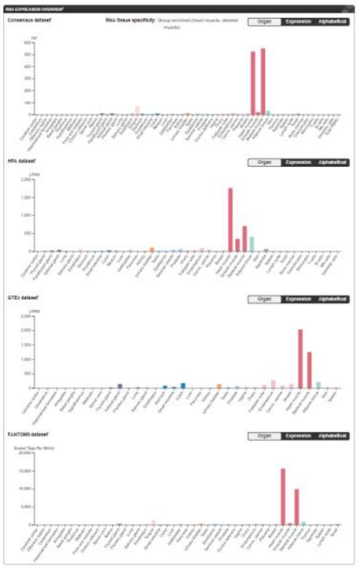

### HSPB7 – Cell type expression in humans

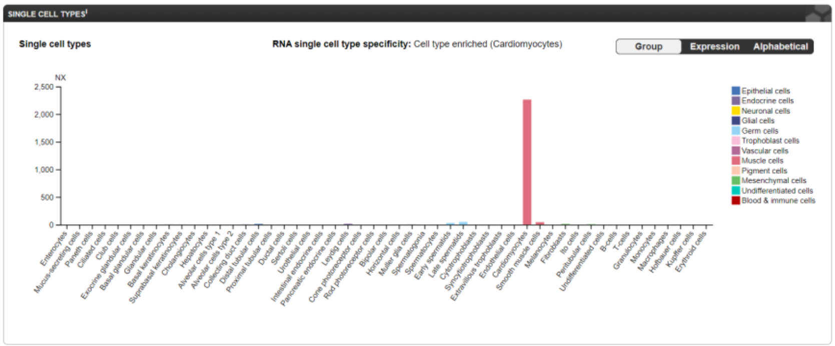


### HSPB7 – Cellular expression in human heart

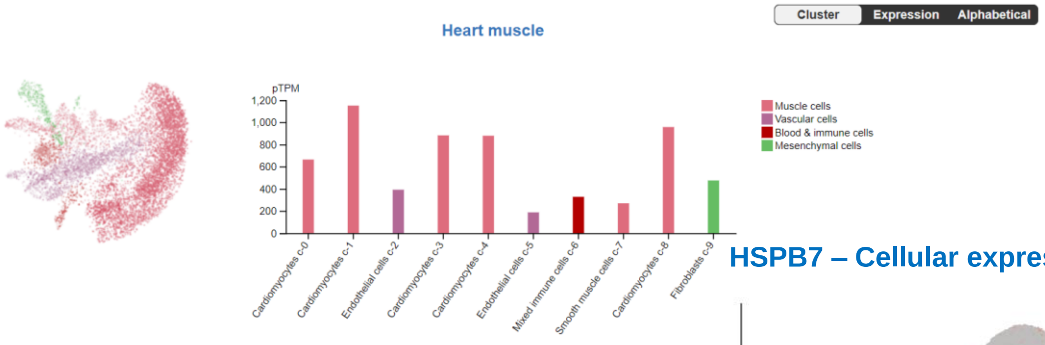


### HSPB7 – Cellular expression in human heart

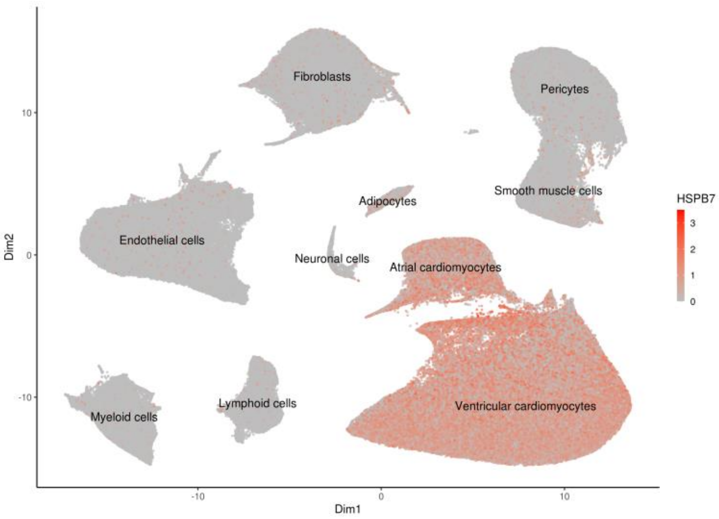


**Fig. S3B. Transcriptomics data analysis on *HSPB7*.** Data shown are the tissue and single cell type mRNA expression levels based on RNA sequencing data sets from GTExPortal (https://gtexportal.org/home/), the Human Protein Atlas (https://www.proteinatlas.org/) and the Heart Cell Atlas (https://www.heartcellatlas.org/).

### LMO2 – Tissue distribution in humans

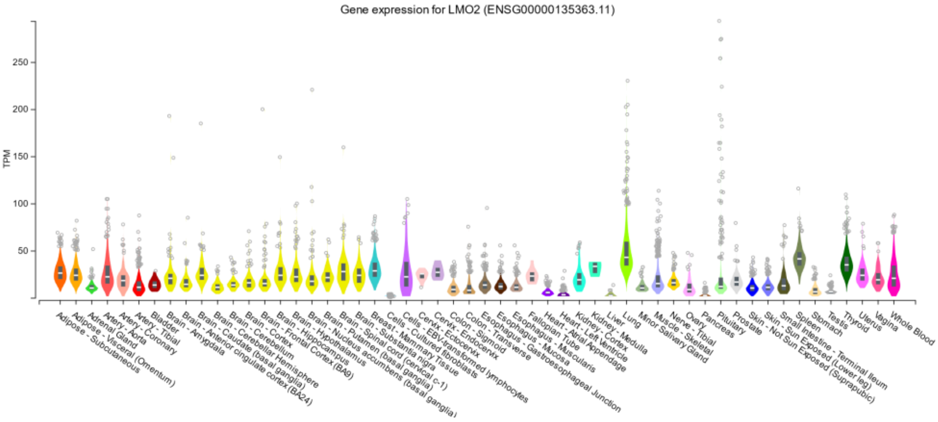


### LMO2 – Tissue distribution in humans

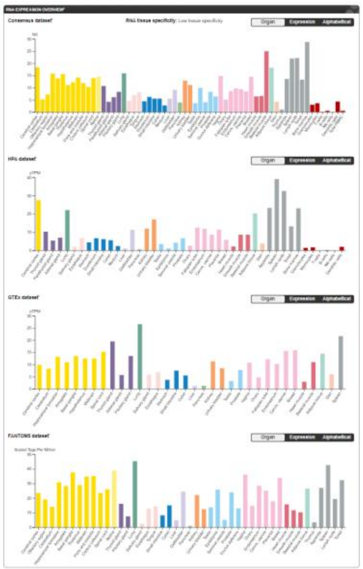

### LMO2 – Cell type expression in humans

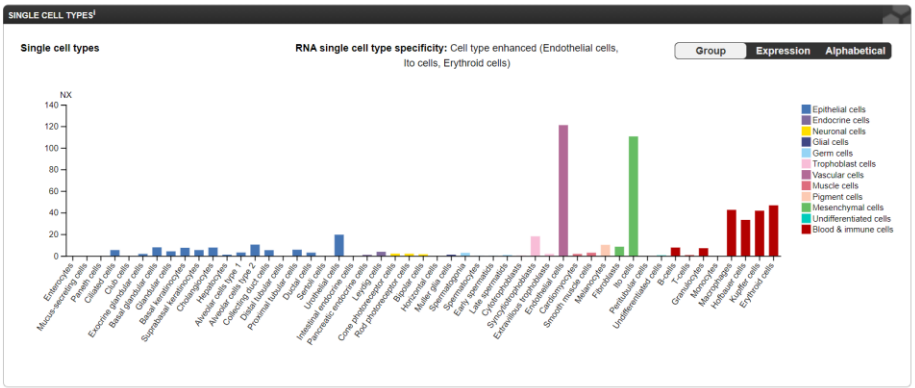


### LMO2 – Cellular expression in human heart

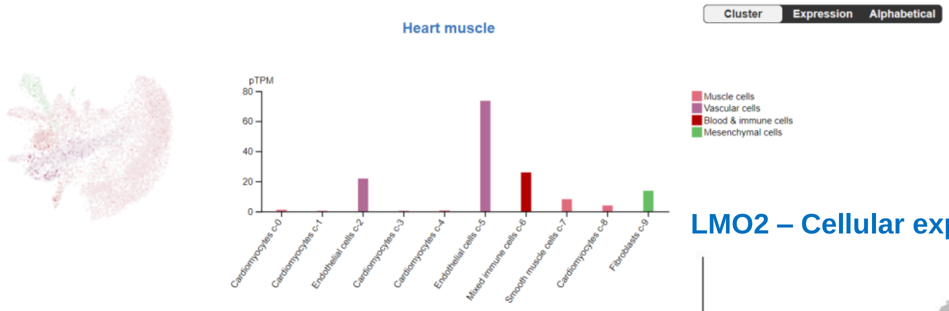


### LMO2 – Cellular expression in human heart

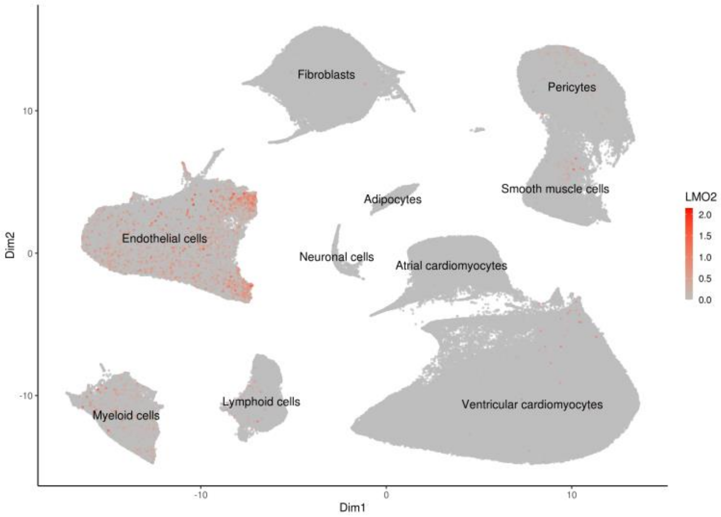


**Fig. S3C. Transcriptomics data analysis on *LMO2*.** Data shown are the tissue and single cell type mRNA expression levels based on RNA sequencing data sets from GTExPortal (https://gtexportal.org/home/), the Human Protein Atlas (https://www.proteinatlas.org/) and the Heart Cell Atlas (https://www.heartcellatlas.org/).

### GATA5 – Tissue distribution in humans

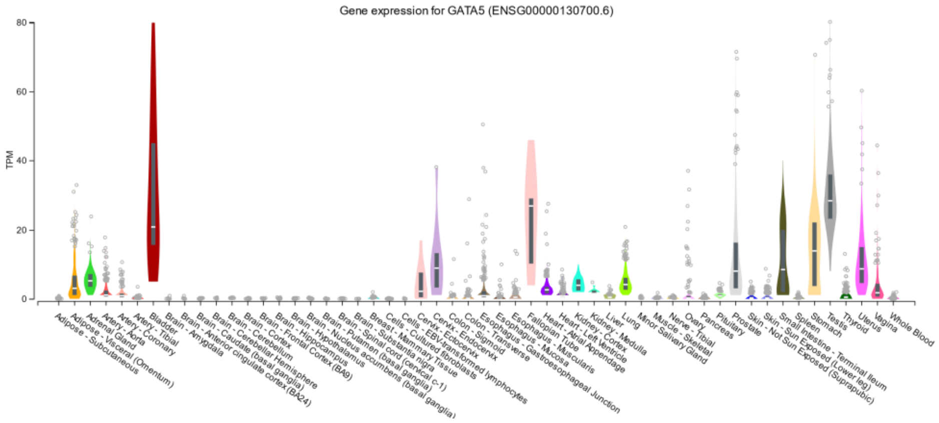


### GATA5 – Tissue distribution in humans

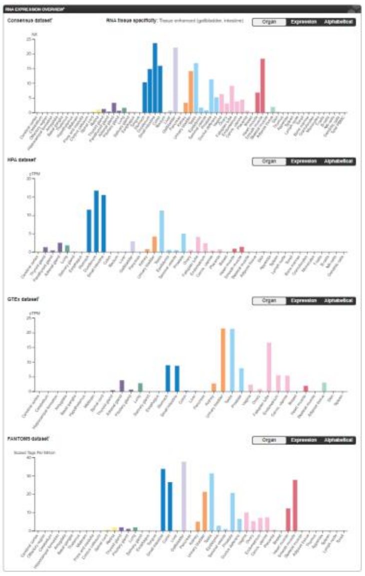

### GATA5 – Cell type expression in humans

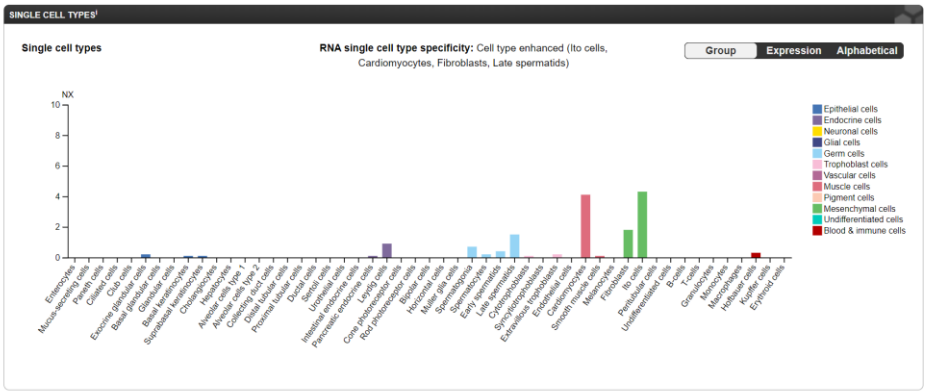


### GATA5 – Cellular expression in human heart

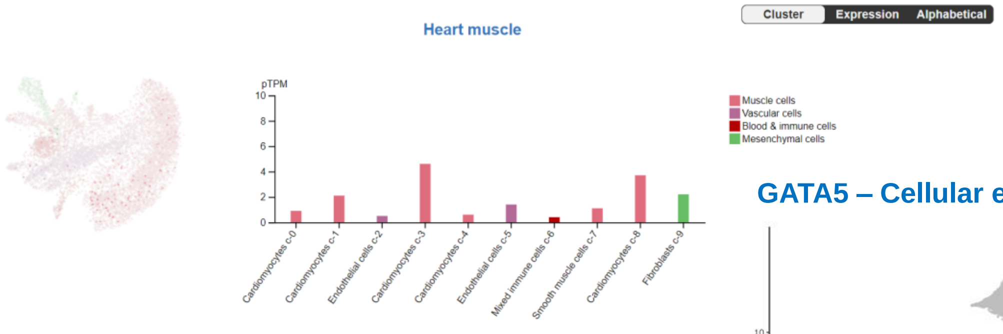


### GATA5 – Cellular expression in human heart

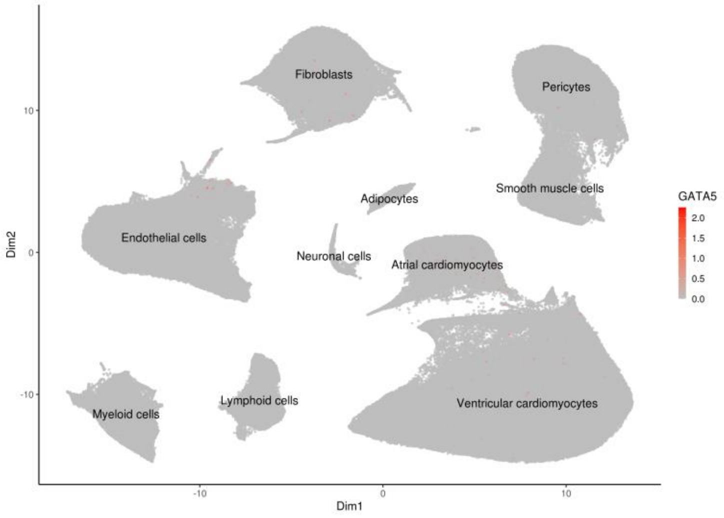


**Fig. S3D. Transcriptomics data analysis on *GATA5*. Data shown are the** tissue and single cell type mRNA expression levels based on RNA sequencing data sets from GTExPortal (https://gtexportal.org/home/), the Human Protein Atlas (https://www.proteinatlas.org/) and the Heart Cell Atlas (https://www.heartcellatlas.org/).

**A)**

| | Malformations at 2 days post fertilisation (2dpf) | | | | | | | | | | | |
|---|---|---|---|---|---|---|---|---|---|---|---|---|
| | n-number (scored) | % Lethality (by 2dpf) | % Larvae Abnormal | Body Shape | Somites | Notochord | Tail | Heart | Face | Neural Tube | Arches/Jaws | Abdomen distended/yolk oedema |
| Cas9 | 110 | 0% | 3% | 0% | 2% | 2% | 0% | 3% | 2% | 2% | 1% | 1% |
| *gata5* g#1 + Cas9 | 95 | 0% | 68% | 0% | 0% | 0% | 0% | 68% | 0% | 0% | 0% | 65% |
| *gata5* g#2 + Cas9 | 57 | 0% | 18% | 0% | 0% | 0% | 0% | 18% | 0% | 0% | 0% | 7% |
| *gata5* g#3 + Cas9 | 60 | 0% | 18% | 0% | 0% | 0% | 0% | 18% | 0% | 0% | 0% | 17% |
| *gata5* g#1,2,3 +Cas9 | 102 | 2% | 87% | 0% | 3% | 1% | 0% | 87% | 3% | 1% | 0% | 94% |

**Aii**

| | Heart-specific malformations at 2dpf | | | | | | |
|---|---|---|---|---|---|---|---|
| | n-number (scored) | % Abnormal | Slow Heartbeat | No Heartbeat | Pericardial Sac-Swollen | Chambers not well defined | Misshapen |
| Cas9 | 110 | 3% | 0% | 0% | 3% | 1% | 2% |
| *gata5* g#1 + Cas9 | 97 | 54% | 3% | 10% | 31% | 2% | 37% |
| *gata5* g#2 + Cas9 | 57 | 18% | 0% | 0% | 18% | 0% | 0% |
| *gata5* g#3 + Cas9 | 60 | 18% | 0% | 0% | 17% | 0% | 3% |
| *gata5* g#1,2,3 +Cas9 | 100 | 87% | 19% | 8% | 73% | 10% | 53% |

**Aiii**

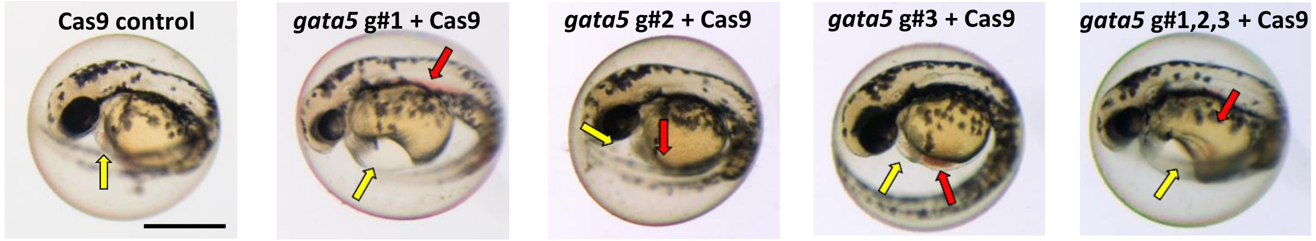


**B)**

| | Malformations at 2 days post fertilisation (2dpf) | | | | | | | | | | | |
|---|---|---|---|---|---|---|---|---|---|---|---|---|
| | n-number (scored) | % Lethality (by 2dpf) | % Larvae Abnormal | Body Shape | Somites | Notochord | Tail | Heart | Face | Neural Tube | Arches/Jaws | Abdomen distended/yolk oedema |
| Cas9 | 42 | 2% | 0% | 0% | 0% | 0% | 0% | 0% | 0% | 0% | 0% | 0% |
| *api5* g#1 + Cas9 | 44 | 0% | 0% | 0% | 0% | 0% | 0% | 0% | 0% | 0% | 0% | 0% |
| *api5* g#2 + Cas9 | 53 | 19% | 0% | 0% | 0% | 0% | 0% | 0% | 0% | 0% | 0% | 0% |
| *api5* g#3 + Cas9 | 44 | 2% | 2% | 0% | 2% | 2% | 0% | 2% | 2% | 2% | 2% | 2% |
| *api5* g#1,2,3 + Cas9 | 50 | 14% | 0% | 0% | 0% | 0% | 0% | 0% | 0% | 0% | 0% | 0% |

**Bii**

| | Heart-specific malformations at 2dpf | | | | | | |
|---|---|---|---|---|---|---|---|
| | n-number (scored) | % Abnormal | Slow Heartbeat | No Heartbeat | Pericardial Sac-Swollen | Chambers not well defined | Misshapen |
| Cas9 | 44 | 0% | 0% | 0% | 0% | 0% | 0% |
| *api5* g#1 + Cas9 | 43 | 0% | 0% | 0% | 0% | 0% | 0% |
| *api5* g#2 + Cas9 | 43 | 0% | 0% | 0% | 0% | 0% | 0% |
| *api5* g#3 + Cas9 | 43 | 2% | 0% | 2% | 0% | 2% | 2% |
| *api5* g#1,2,3 + Cas9 | 41 | 0% | 0% | 0% | 0% | 0% | 0% |

**Biii**

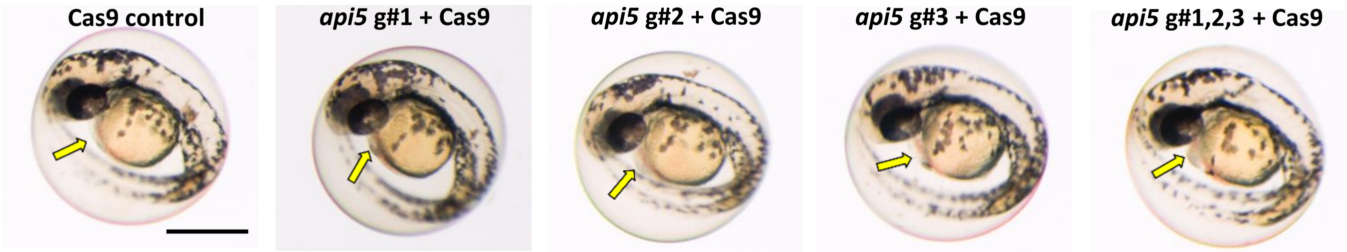


**Fig. S4A. Comparison of morphological endpoints measured in 2dpf zebrafish. Panel A)** General whole body morphological endpoints measured following injection of Cas9 alone, or injection of *gata5* gRNAs 1, 2 and 3 in isolation or combined (g#1, g#2, g#3 or g#1,2,3 respectively) all with Cas9. Data are shown as the % incidence of abnormalities under each of the morphological categories. **Ai**: Expansion of the heart-specific endpoints measured showing the full range of endpoints scored within this category, across the same treatments. **Aii**: Example images of typical 2dpf animals within each treatment category. The yellow arrows show the position of the pericardial membrane and the extent of pericardial oedema in that animal, and the red arrows shows regions of blood pooling; **Panel B)** As Panel A, but for *api5*. More detailed whole body morphological endpoint scoring and cardiovascular functional assessment was undertaken on the two guide combinations showing the strongest phenotypes from the 2dpf data. Scale bar shown in the first image of each panel represents 500µm.

**C)**

| Malformations at 2 days post fertilisation (2dpf) | n-number (scored) | % Lethality (by 2dpf) | % Larvae Abnormal | Body Shape | Somites | Notochord | Tail | Heart | Face | Neural Tube | Arches/Jaws | Abdomen distended/yolk oedema |
|---|---|---|---|---|---|---|---|---|---|---|---|---|
| Cas9 | 48 | 0% | 0% | 0% | 0% | 0% | 0% | 0% | 0% | 0% | 0% | 0% |
| *hspb7* g#1 + Cas9 | 47 | 0% | 2% | 0% | 0% | 0% | 0% | 2% | 0% | 0% | 0% | 0% |
| *hspb7* g#2 + Cas9 | 48 | 0% | 2% | 2% | 2% | 2% | 2% | 2% | 2% | 2% | 2% | 0% |
| *hspb7* g#3 + Cas9 | 41 | 0% | 12% | 0% | 10% | 7% | 5% | 10% | 10% | 10% | 10% | 10% |
| *hspb7* g#1,2,3 + Cas9 | 48 | 0% | 17% | 10% | 13% | 10% | 13% | 10% | 13% | 13% | 8% | 13% |

**Cii**

| Heart-specific malformations at 2dpf | n-number (scored) | % Abnormal | Slow Heartbeat | No Heartbeat | Pericardial Sac Swollen | Chambers not well defined | Misshapen |
|---|---|---|---|---|---|---|---|
| Cas9 | 48 | 0% | 0% | 0% | 0% | 0% | 0% |
| *hspb7* g#1 + Cas9 | 47 | 2% | 0% | 0% | 2% | 0% | 0% |
| *hspb7* g#2 + Cas9 | 48 | 2% | 0% | 2% | 0% | 2% | 2% |
| *hspb7* g#3 + Cas9 | 41 | 10% | 5% | 0% | 5% | 0% | 5% |
| *hspb7* g#1,2,3 + Cas9 | 48 | 10% | 0% | 4% | 10% | 6% | 8% |

**Ciii**

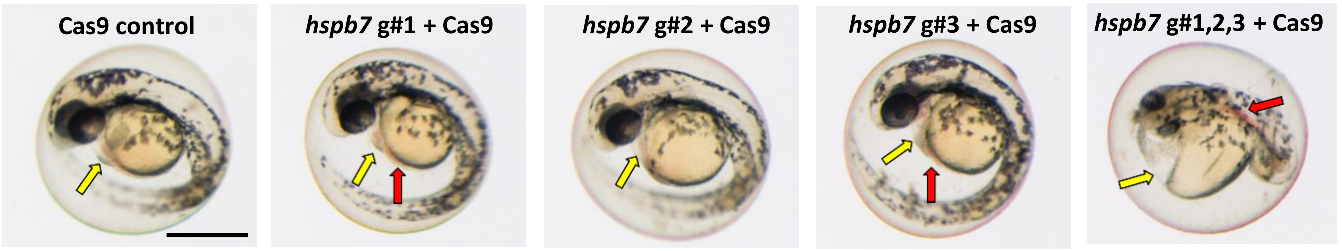


**D)**

| Malformations at 2 days post fertilisation (2dpf) | n-number (scored) | % Lethality (by 2dpf) | % Larvae Abnormal | Body Shape | Somites | Notochord | Tail | Heart | Face | Neural Tube | Arches/Jaws | Abdomen distended/yolk oedema |
|---|---|---|---|---|---|---|---|---|---|---|---|---|
| Cas9 | 46 | 8% | 7% | 0% | 2% | 2% | 2% | 7% | 2% | 2% | 2% | 2% |
| *lmo2* g#1 + Cas9 | 3 | 94% | 33% | 0% | 33% | 33% | 33% | 33% | 33% | 33% | 33% | 33% |
| *lmo2* g#2 + Cas9 | 45 | 10% | 91% | 2% | 20% | 7% | 11% | 91% | 24% | 11% | 9% | 18% |
| *lmo2* g#3 + Cas9 | 43 | 24% | 91% | 5% | 12% | 5% | 5% | 91% | 14% | 9% | 7% | 44% |
| *lmo2* g#1,2,3 + Cas9 | 12 | 76% | 92% | 25% | 67% | 50% | 58% | 92% | 58% | 58% | 42% | 58% |

**Dii**

| Heart-specific malformations at 2dpf | n-number (scored) | % Abnormal | Slow Heartbeat | No Heartbeat | Pericardial Sac Swollen | Chambers not well defined | Misshapen |
|---|---|---|---|---|---|---|---|
| Cas9 | 46 | 7% | 2% | 0% | 7% | 0% | 4% |
| *lmo2* g#1 + Cas9 | 3 | 33% | 0% | 33% | 33% | 33% | 33% |
| *lmo2* g#2 + Cas9 | 45 | 91% | 7% | 7% | 73% | 11% | 82% |
| *lmo2* g#3 + Cas9 | 43 | 91% | 2% | 2% | 60% | 2% | 81% |
| *lmo2* g#1,2,3 + Cas9 | 12 | 92% | 8% | 17% | 58% | 25% | 92% |

**Diii**

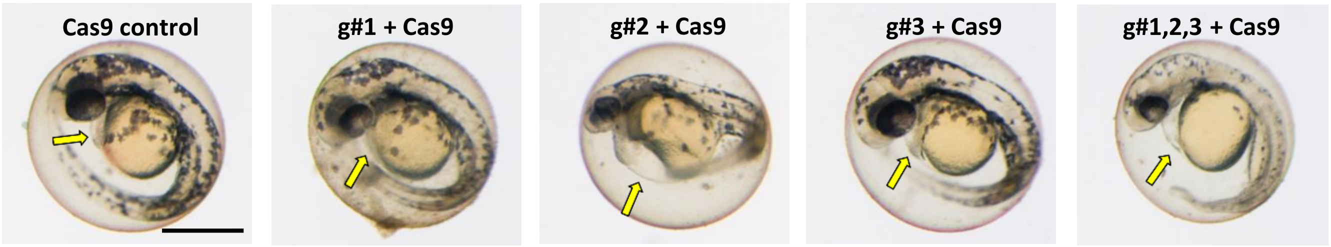


**Fig. S4B. Comparison of morphological endpoints measured in 2dpf animals. Panel C)** General whole body morphological endpoints measured following injection of Cas9 alone, or injection of *hspb7* gRNAs 1, 2 and 3 in isolation or combined (g#1, g#2, g#3 or g#1,2,3 respectively) all with Cas9. Data are shown as the % incidence of abnormalities under each of the morphological categories. **Ci**: Expansion of the heart-specific endpoints measured showing the full range of endpoints scored within this category, across the same treatments. **Cii**: Example images of typical 2dpf animals within each treatment category. The yellow arrows show the position of the pericardial membrane and the extent of pericardial oedema in that animal, and the red arrows shows regions of blood pooling; **Panel D)** as Panel C) but for *lmo2*. More detailed whole body morphological endpoint scoring and cardiovascular functional assessment was undertaken on the two guide combinations showing the strongest phenotypes from the 2dpf data. Scale bar shown in the first image of each panel represents 500µm.

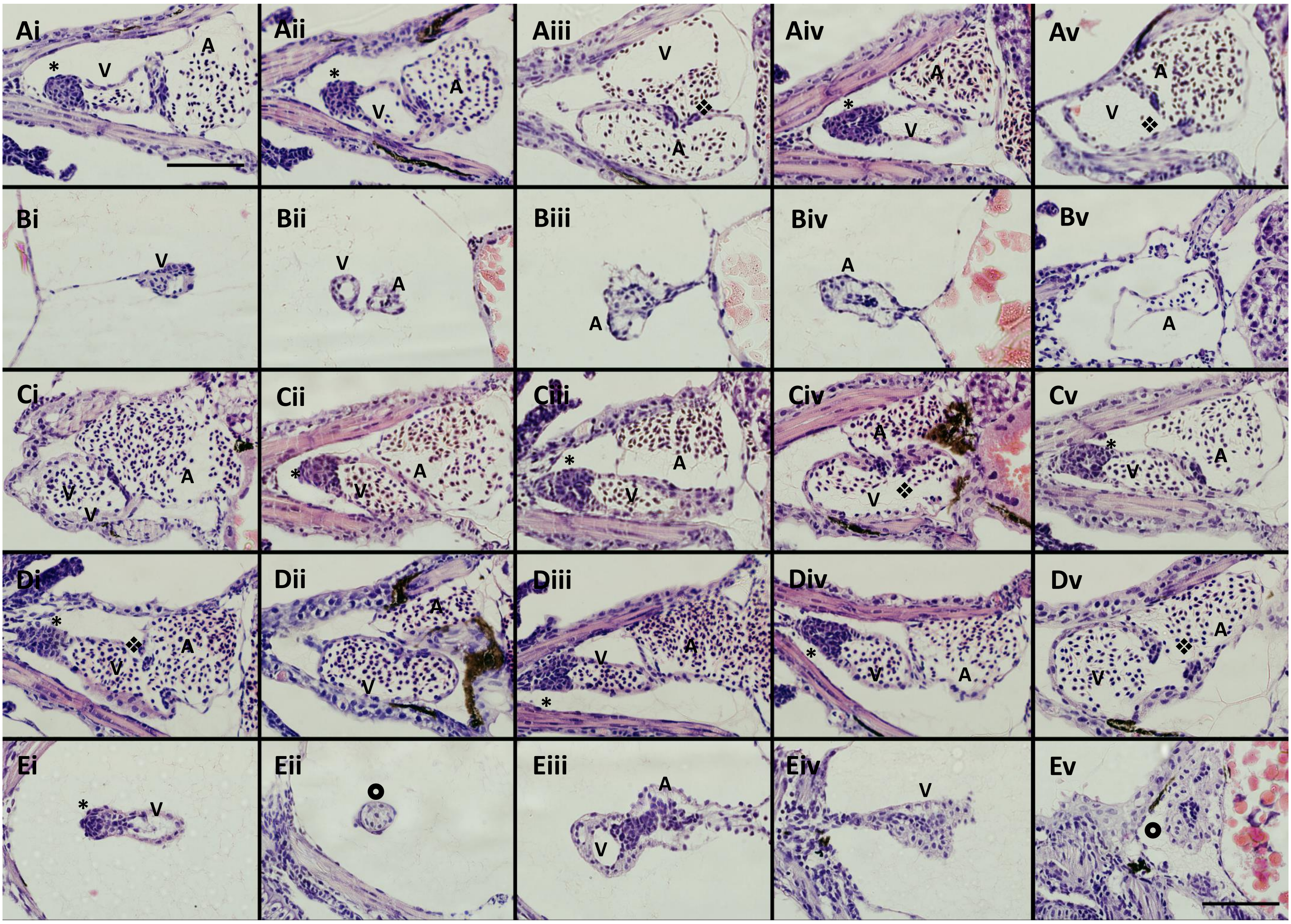


**Fig. S5. Haematoxylin and eosin stained coronal sections through the hearts of 4dpf CRISPR-mutant zebrafish.** In each row are sections from 5 animals (i-v) subjected to A) Cas9 injection; B) gata5 g#1,2,3 + Cas9 injection; C) *api5* g#1,2,3 + Cas9 injection; D) *hspb7* g#1,2,3 + Cas9 injection and E) *lmo2* g#1,2,3 + Cas9 injection. In each panel, animals are orientated with the head to the left, and viewed in the dorsal plane at a magnification of 40x (Scale bar shown in top left and bottom right images represents 200µm). On various panels, labelled structures include the ventricle (v); atrium (A); bulbus arteriosus (*); atrioventricular value (❖); and unspecified heart tissue (⦿).

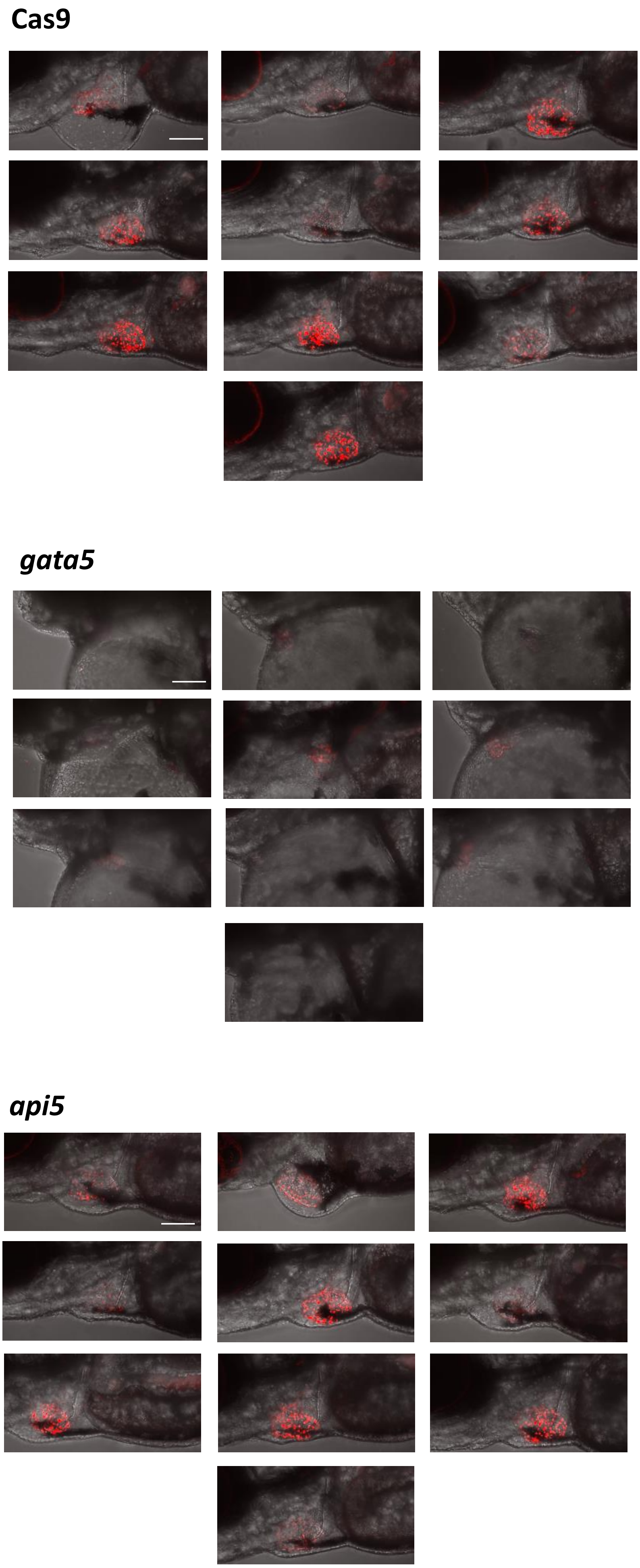


**Fig. S6A. Confocal maximum intensity projection images of hearts from *cmlc2::DsRed2-nuc* mutant larvae**. The images shown are the combined transmitted light and *cmlc2::DsRed2-nuc* fluorescence signals from 10 randomly sampled larvae per treatment, with cardiomyocytes fluorescing in red, especially prominently in the ventricle (all laser settings identical). The top set of images shows hearts from Cas9 injected (injection controls) and then from embryos injected with *gata5* g#1,2,3 + Cas9 (middle), and *api5* g#1,2,3 + Cas9, (bottom). Scale bar shown in the first image of each panel represents 100 µm.

**Cas9**

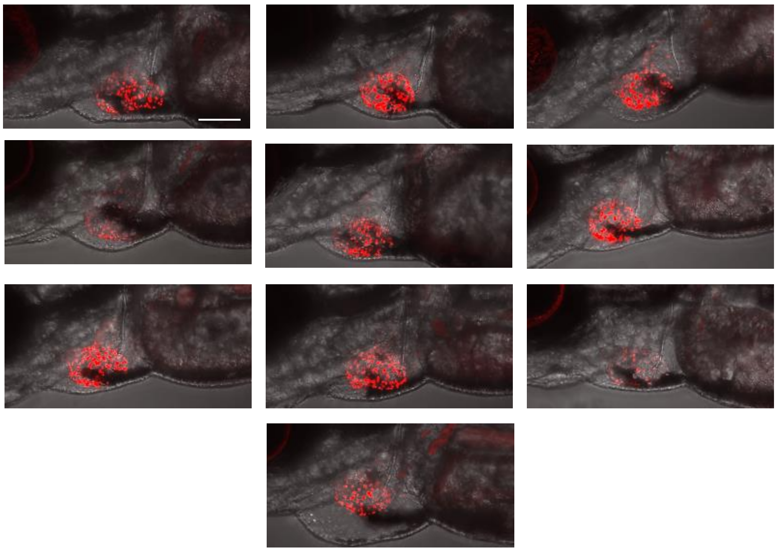

***hspb7***

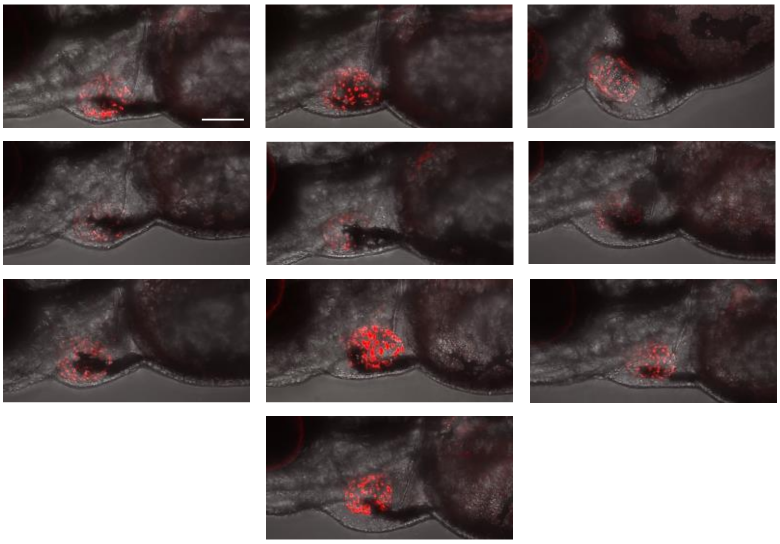

***lmo2***

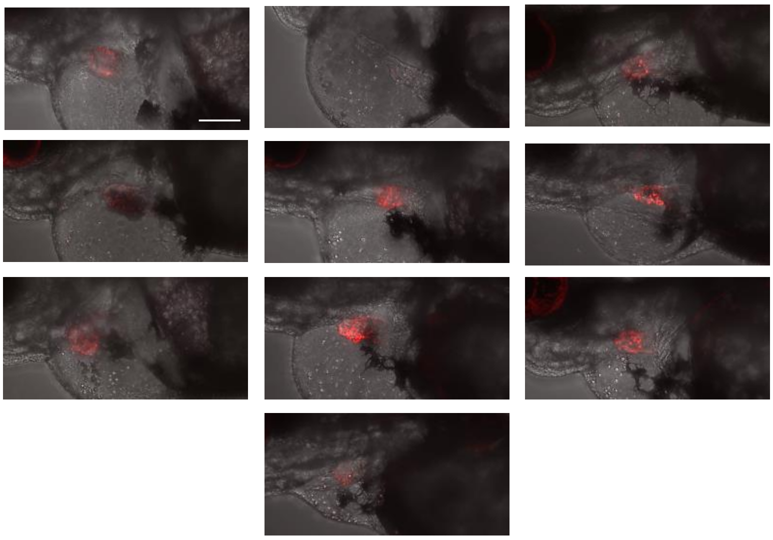

**Fig. S6B. Confocal maximum intensity projection images of hearts from *cmlc2::DsRed2-nuc* mutant larvae**. The images shown are the combined transmitted light and *cmlc2::DsRed2-nuc* fluorescence signals from 10 randomly sampled larvae per treatment, with cardiomyocytes fluorescing in red, especially prominently in the ventricle (all laser settings identical). The top set of images shows hearts from Cas9 injected (injection controls) and then from embryos injected with *hspb7* g#1,2,3 + Cas9 (middle) and *lmo2* g#2 + Cas9 (bottom). Scale bar shown in the first image of each panel represents 100 μm.

### SUPPLEMENTARY VIDEOS

**A)**

| **Cas9** | ***gata5* g#1 + Cas9** | ***gata5* g#1,2,3 + Cas9** |
|---|---|---|
| F3 cas9_BF_0001.mpg | F7 Guide 1_BF_0001.mpg | F8 Guide 1 2 3 combined_BF_0001.mpg |
| F3 cas9_HB_0001.mpg | F7 Guide 1_HB_0001.mpg | F8 Guide 1 2 3 combined_HB_0001.mpg |

**B)**

| **Cas9** | ***api5* g#1 + Cas9** | ***api5* g#1,2,3 + Cas9** |
|---|---|---|
| Cas9 F1_BF_0001.mpg | Guide 1 F8_BF_0001.mpg | Combined 1 2 3 F5_BF_0001.mpg |
| Cas9 F1_HB_0001.mpg | Guide 1 F8_HB_0001.mpg | Combined 1 2 3 F5_HB_0001.mpg |

**C)**

| **Cas9** | ***hspb7* g#3 + Cas9** | ***hspb7* g#1,2,3 + Cas9** |
|---|---|---|
| Cas9 F5_BF_0001.mpg | Guide 3 Hspb7 F6_BF_0001.mpg | Combined Hspb7 F1 REPEATED_BF_0001.mpg |
| Cas9 F5_HB_0001.mpg | Guide 3 Hspb7 F6_HB_0001.mpg | Combined Hspb7 F1 REPEATED_HB_0001.mpg |

**D)**

| **Cas9** | ***lmo2* g#1,2,3 + Cas9** | ***lmo2* g#2 + Cas9** |
|---|---|---|
| Cas9 F2_BF_0001.mpg | combined F4 DEFORMED_BF_0001.mpg | Guide 2 F5 DEFORMED_BF_0001.mpg |
| Cas9 F2_HB_0001.mpg | combined F4 DEFORMED_HB_0001.mpg | Guide 2 F5 DEFORMED_HB_0001.mpg |

**Videos of cardiovascular function to accompany Figure 5 in the main manuscript. Panel A)** Videos showing the vasculature in the upper row, and the heart recorded from example fish injected with Cas9 alone (left), *gata5* g#1 + Cas9 (middle) and *gata5* g#1,2,3 + Cas9 (right). **Panel B)** As Panel A), for Cas9 alone (left), *api5* g#1 + Cas9 (middle) and *api5* g#1,2,3 + Cas9 (right). **Panel C)** As Panel A), for Cas9 alone (left), *hspb7* g#3 + Cas9 (middle) and *api5* g#1,2,3 + Cas9 (right). **Panel D)** As Panel A), for Cas9 alone (left), *lmo2* g#1,2,3 + Cas9 (middle) and *api5* g#2 + Cas9 (right).

### SUPPLEMENTARY DATASETS

**Data S1. Data S1_Network Analysis.xlsx**

**Data S2. Data S2_Differential Expression.xlsx**

**Data S3. Data S3_Genetic Associations.xlsx**

**Data S4. Data S4_Homologies.xlsx**
